# Encoding of an engram for food location by satiety-promoting Drd2 hippocampal neurons

**DOI:** 10.1101/439349

**Authors:** Estefania P. Azevedo, Lisa Pomeranz, Jia Cheng, Marc Schneeberger, Sarah Stern, Katherine Doerig, Paul Greengard, Jeffrey M. Friedman

**Author notes:** Correspondence should be addressed to, Jeffrey M. Friedman.

## Abstract

Associative learning guides feeding behavior in mammals in part by using cues that link location in space to food availability. However, the elements of the top-down circuitry encoding the memory of the location of food is largely unknown, as are the high-order processes that control satiety. Here we report that hippocampal dopamine 2 receptor (D2R) neurons are specifically activated by food and that modulation of their activity reduce food intake in mice. We also found that activation of these neurons interferes with the valence of food and the acquisition of a spatial memory linking food to a location via projections from the hippocampus to the lateral septum. Finally, we showed that inputs from lateral entorhinal cortex (LEC) to the hippocampus can also drive satiety via activation of D2R cells. These data describe a previously unidentified function for hippocampal D2R cells to regulate feeding behavior and identifies a LEC->Hippocampus->Septal high-order circuit that encodes the memory of food location.

## INTRODUCTION

Homeostatic control of energy balance is vital for the survival of all animals. Specific populations of neurons in the brain control food intake and energy balance by integrating a panoply of relevant sensory and hormonal signals, in turn orchestrating an adaptive behavioral response (Waterson and Horvath, 2015). Defined neural populations in several brain regions, including the hypothalamus, midbrain reward centers (Domingos et al., 2011), parabrachial nucleus, amygdala, pre-frontal cortex, dorsal raphe nucleus and others, have all been shown to control appetite (Aponte et al., 2010; Holland, 2004; Nectow et al., 2017; Waterson and Horvath, 2015) and mutations in genes that impact the function of these neuronal populations can be associated with obesity or anorexia (Herman et al., 2016; Mutch and Clément, 2006; Nectow et al., 2017). Thus, the neural processes by which these and other brain regions integrate relevant environmental cues to regulate feeding play a critical role in enabling an animal to efficiently obtain food. However, while it is especially important for an animal to link a memory of a specific location in space to the availability of food, the cellular mechanisms responsible for this association are largely unknown.

The role of the hippocampus in episodic memory and spatial location in rodents and humans is well established and lesions in this brain region can result in memory loss (Eichenbaum, 2000). The hippocampus is divided into discrete anatomic regions that control diverse processes such as location in space and time and the memory of an unpleasant (aversive) experience (Eichenbaum, 2014; Hartley et al., 2013). Recently, several groups have also shown that hippocampal dysfunction can alter feeding behavior (Kanoski and Grill, 2015). For example, impairments in a variety of complex mental representations due to poor memory (Hussey et al., 2012) and deficits in the regulation of satiety have been reported in patients diagnosed with retrograde amnesia (Higgs et al., 2011; Rozin et al., 1998). In addition, it has been shown that imagining food can induce physiological changes, such as the production of gastric juices and salivation (Nederkoorn and Jansen, 2001) and that this imagined consumption can decrease food intake in humans through habituation (Morewedge et al., 2010). This suggests that cognitive processes can also regulate satiety, and potentially even (partially) explain why some patients can maintain weight loss for long periods of time, while most patients cannot (Brockmeyer et al., 2017). However, the neural mechanisms responsible for the modulation of feeding by cortical structures are poorly understood as are the role of specific hippocampal neural populations in these processes. Thus, the molecular identification of a specific ensemble in the hippocampus that responds to food would enable a functional analysis of a top-down circuit that processes sensory information including spatial cues to modulate energy intake.

The hippocampus is also known to encode an engram of high valence, contextual experiences including footshocks received in a specific context (Ramirez et al., 2013). As termed by Semon more than 100 years ago, an engram refers to a (presumed) physical, biological alteration in the brain that occurs after a specific experience, thus encoding a memory of that experience (Josselyn et al., 2015). Semon suggested that changes in the activity or other aspects of the function of specific neuronal populations after high valence experiences leads to the formation of a memory of that experience (engram).

An animal’s ability to associate food with a particular location is critical for efficiently finding food and conserving energy. We hypothesized that the hippocampus might also respond to food similarly to the way it responds to cues associated with other experiences (Eichenbaum, 2014; Sanders et al., 2015). We thus set out to determine whether the presentation of a meal could activate specific hippocampal cells, so we could then evaluate whether those neurons play a role in the formation of an engram for food, to in turn regulate hunger or satiety. Here, we report that dopamine 2 receptor (D2R) neurons in the hilar region of the hippocampus are activated by food. The selective modulation of the activity of D2R neurons is sufficient to induce changes in food intake and negatively regulate the acquisition of a memory for the location of food. Furthermore, specific projections from these neurons to the septal area control feeding and avoidance behavior while a projection of glutamatergic neurons in lateral entorhinal cortex (LEC) can regulate feeding by activating hippocampal D2R cells. These findings describe a previously unidentified role for D2R cells within the hippocampus and define some of the elements of a multimodal neural LEC->Hippocampus->Septal circuit that regulates food intake and conveys a memory of spatial cues associated with food location.

## RESULTS

### Hippocampal cells respond to energy states and control feeding

To determine whether hippocampal neurons show a change in activity after food deprivation, we assayed the levels of the immediate early gene c-fos in the hippocampus of fasted and fed mice (Figure 1A and 1B). After an overnight fast, several hippocampal areas, including CA1, CA3, dentate gyrus (DG) and the hilus, showed significantly lower levels of c-fos immunoreactivity compared to fed mice (Figure 1B). Based on this preliminary finding, we next evaluated whether manipulating the activity of hippocampal cells in mice could influence food intake (Figure 1C-E). We targeted the inhibitory Gi(hM4Di) or excitatory Gq-coupled (hM3Dq) designer receptor exclusively activated by designer drugs (DREADDs) to glutamatergic neurons in the hippocampus by injecting viruses expressing these constructs under the control of the CamkIIa promoter in wild-type mice (Figure 1C). As controls, we used mice bilaterally injected with a mCherry-expressing virus in the same brain region (Figure 1C; right panel). We found that a single CNO injection significantly increased food intake for 24h in mice expressing the hM4Di inhibitory DREADD (Figure 1D, *Acute* and Figure S1B). In contrast, acute activation of these neurons expressing the activating hM3Dq DREADD by CNO acutely decreased food intake (Figure 1D, right panel, *Acute* and Figure S1C). Chronic treatment with CNO showed quantitatively greater effects and after three days, food intake was significantly increased after inhibiting glutamatergic dorsal neurons in the hippocampus compared to controls (Figure 1E, left panel, *Chronic*), while chronic activation led to hypophagia (Figure 1E, right panel, *Chronic*). Saline injections in DREADD-expressing mice as well as CNO injection in mCherry controls did not alter food intake (Figure 1D and 1E). These data suggest that a glutamatergic subpopulation in the dorsal hippocampus can modulate food intake in mice.

**Figure 1.**
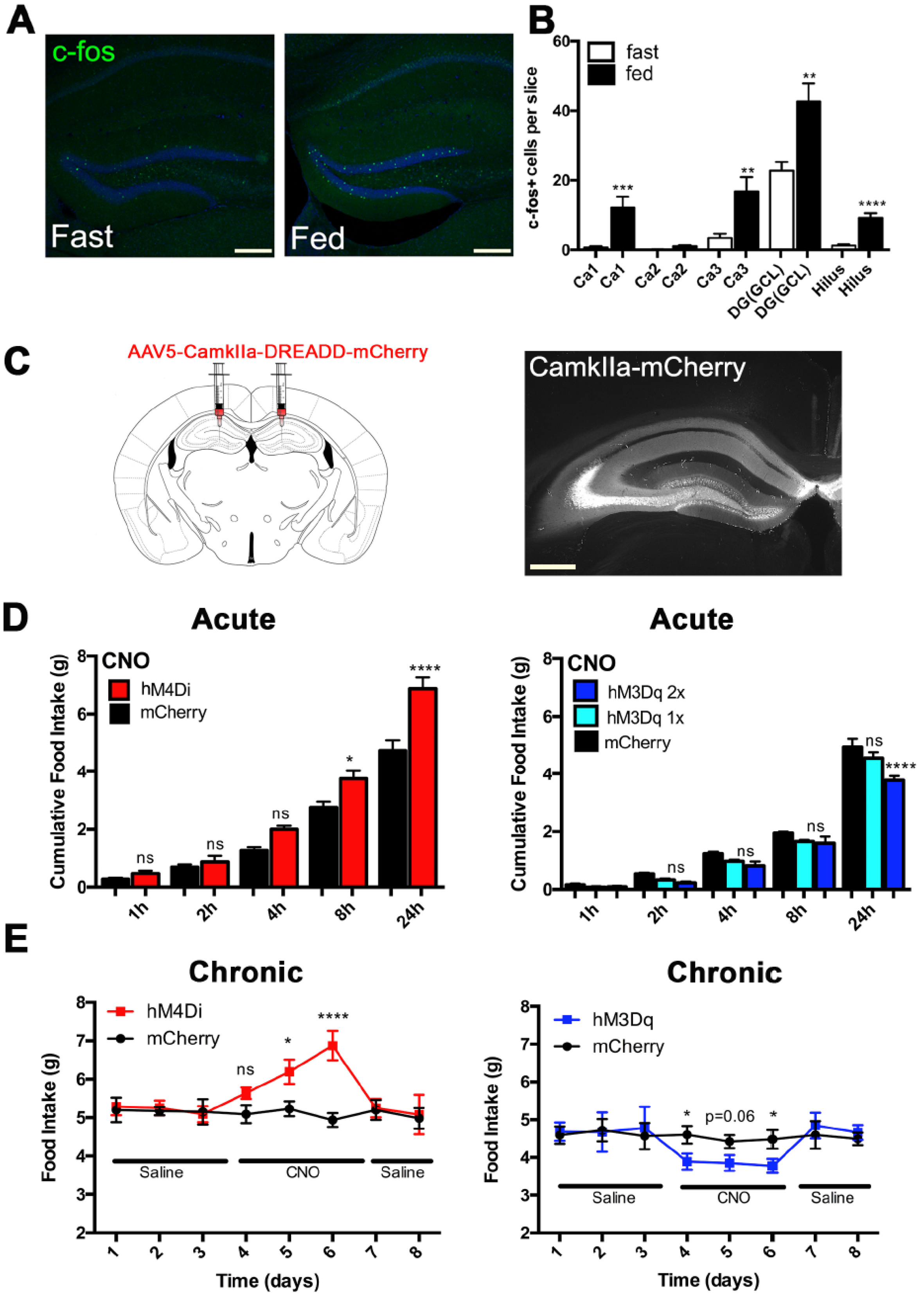
Hippocampal cells respond to nutritional changes and control food intake. (A and B) Analysis of c-fos expression (green) in (A-left) fast and (A-right) fed mice. Scale, 200 *μ*m. The number of c-fos+ cells per slice was quantified and plotted for fast and fed groups (n=3/group). (C) Schematic (left panel) and representative (right panel) of AAV5-CamkIIa-DREADD(hM3Dq and hM4Di)-mCherry expression in the dorsal hippocampus of wild-type mice. Scale, 400 *μ*m. (D) Left panel; acute cumulative food intake (g) after a single injection of CNO (i.p; 1 mg/kg) in hM4Di-injected mice (red bars; n=8) and mCherry-injected mice (black bars; n=8). One-way ANOVA with Bonferroni correction, *p<0.05 and ****p<0.0001. Right Panel; acute cumulative food intake (g) after a single (light blue bars) or double (dark blue bars) injection of CNO (i.p; 1 mg/kg) in hM3Dq-injected mice (light and dark blue bars; n=8) and mCherry-injected mice (black bars; n=8). One-way ANOVA with Bonferroni correction, ****p<0.0001. (E) Left panel; chronic food intake (g) after 3 days of saline injection, 3 days of CNO injection (i.p; 1mg/kg), 2 days of saline injection in hM4Di-injected mice (red lines; n=8) and mCherry-injected mice (black lines; n=8). One-way ANOVA with Bonferroni correction, *p<0.05 and ****p<0.0001. Right Panel; chronic food intake (g) after 3 days of saline injection, 3 days of CNO injection (i.p; 1mg/kg, 12/12h), 2 days of saline injection in hM3Dq-injected mice (blue lines; n=9) and mCherry-injected mice (black lines; n=8). One-way ANOVA with Bonferroni correction, *p<0.05.

### A Hippocampal Engram Linking Food to Spatial Location

The hippocampus plays a key role in short-term memory and helps to construct a representation of a memory, or engram (Josselyn et al., 2015). We hypothesized that the appetitive characteristics of food (appearance, taste, odor or calories) and its context or location could create an engram for a meal. We evaluated this possibility by devising a behavioral task in which food is associated with a specific spatial location in a familiar context (Figure 2A). In the novel paradigm we developed, experimental and control groups of mice were exposed to a novel environment for 5 minutes (*Habituation*). They were then returned to their home cage and fasted. After the fast, control mice were returned to the previous environment and empty food cups were placed on opposite sides of the chamber. This group is referred to as *Context*. In the experimental group, mice were fasted overnight, and placed in the environment (to which they were habituated) but in this case, food was added in one of the two food cups in a specific, consistent location while the other remained empty. This group is referred to as *Context+Food*. As expected, fasted mice placed into an environment with the empty food cups (*Context*) spent equal amounts of time in both quadrants containing the cups. In contrast, in the experimental group (*Context+Food*), mice spent a significantly greater amount of time in the quadrant where the foodcontaining cup was placed, resulting in a discrimination index of 0.63±0.1 (Figure 2A; right panel, *Context+Food*, blue bar, p<0.05). Note, because this was conducted for only five minutes, the mice were only able to consume a minimal amount food and thus were not satiated at the end of the training period (<0.01g of food consumed) The above data are shown as a discrimination index; the time spent in each cup-containing quadrant divided by the sum of the time spent in the two quadrants in which the cups were placed. The Discrimination Index (DI) used here has been previously employed (Hattiangady et al., 2014; Ramos-Chavez et al., 2015) and accurately reflects the performance of the task as animals spend little time in the remaining empty quadrants (Figure S2A, white bars).

**Figure 2.**
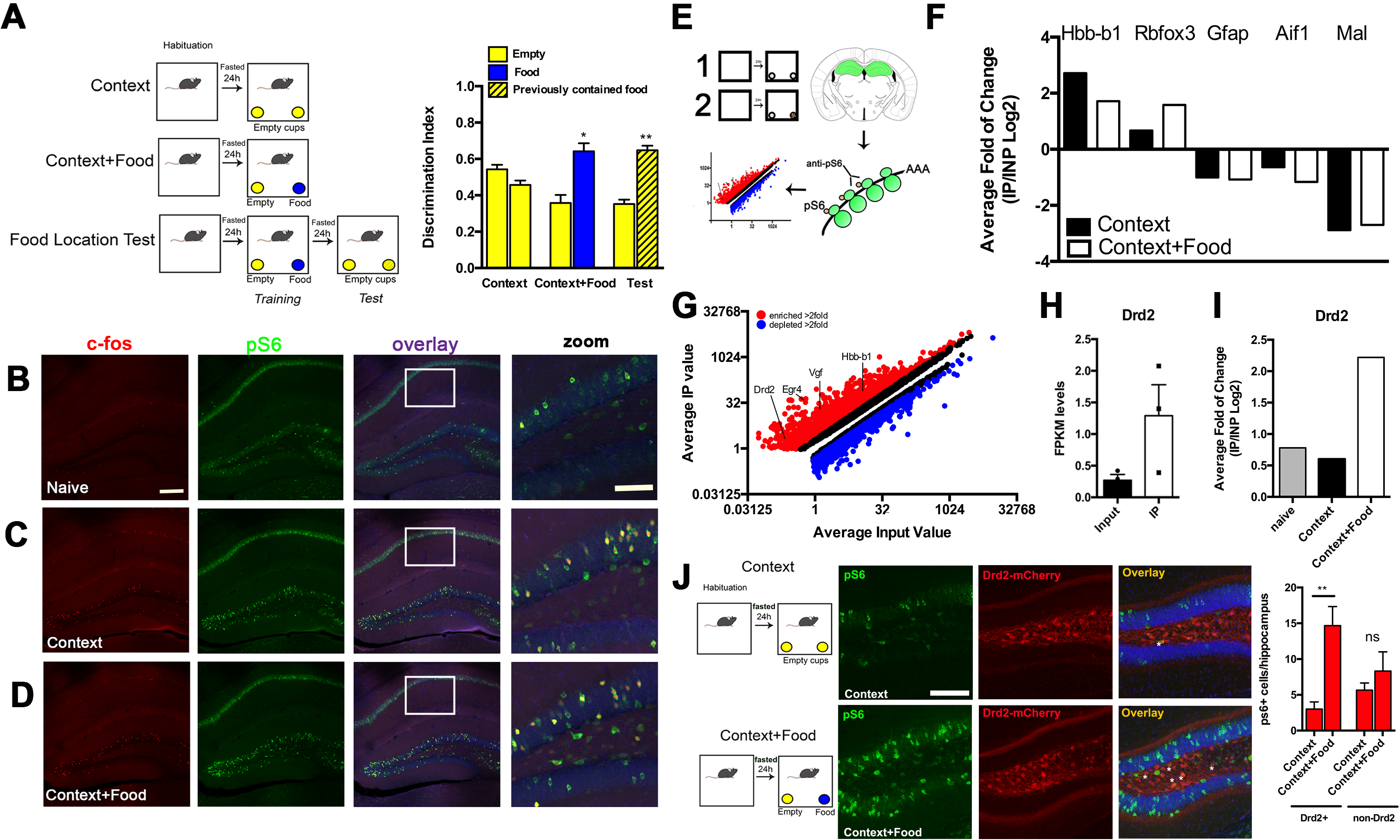
Molecular profiling of hippocampal cells. (A) Left panel, schematic representation of food exposure in a context. Briefly, mice were habituated to an empty arena and fasted overnight. After 24h, mice were exposed to empty food cups (*Context*; yellow) or to one food cup containing mouse chow pellets (blue, *Context+Food*). An additional day (test) was added to the *Context+Food* task enabling us to test memory acquisition and retrieval (Food Location Test). Right panel, discrimination index after 5min exploration time of empty (yellow bars), food containing cups (blue bars) and cups that previously contained food but now were empty (yellow hatched bars). Paired Student’s t-test, ** p<0.01, *p<0.05; n=5. (B-D) Images showing c-fos (red) and pS6 (green) expression in the dorsal hippocampus of (B) naive mice, (C) *Context* mice and (D) *Context+Food* mice. Overlay is shown in the second right panel and overlay zoom of the hilar region is shown in the first right panel. Scale bars, 200 *μ*m and zoom 125 *μ*m. (E) Schematic representation of the molecular profiling of hippocampal cells from *Context* and *Context+Food* groups. (F) Average fold of change (IP/INP Log2) of Hbb-b1, Rbfox3, Gfap, Aif1, Mal genes in *Context* (black bars) and *Context+Food* (white bars) (n=3). Hbb-b1 is used a protocol control (see Methods). (G) Plot depicting the average IP value and average input values (log2) of all genes analyzed. Enriched genes (>2-fold; red) and depleted genes (< 2-fold; blue) are shown and activation or target gene markers are shown: Drd2, Egr4, Vgf and Hbb-b1 (n=3). All genes depicted have q values <0.05 as calculated by Cufflinks. (H) FPKM levels of Drd2 RNA in *Context+Food* groups: input (black bars) and IP (white bars) are compared (n=3). (I) Average fold of change (IP/INP Log2) of Drd2 RNA between naïve (grey bars), *Context* (black bars) and *Context+Food* (white bars): q values are <0.05 for *Context+Food, q=*0.48 for *Context* and q=0.24 for naïve groups. (J) Schematic representation of the exposure of mice to food in a context (left panel). After, brains of Drd2-mCherry mice were stained for pS6 (right panel; green) and Drd2+ and non-Drd2 cells per hippocampus were counted for each group; One-way ANOVA with Bonferroni correction, **p<0.01, n=3. Scale bars, 125 *μ*m.

In order to test whether exposing fasted mice to food in a familiar context was capable of creating an engram, the experimental group was then subjected to an additional day of test (Figure 2A; Food Location Test). After training (i.e. placing fasted animals in a cage for 5 minutes with two food cups, one of which contained food), the animals were fasted for 24h and again placed for five minutes in the familiar chamber but now with two clean, empty cups without food (Test session). The amount of time spent in the quadrant where the food had been previously placed during training was then monitored. We found that during the Test period, mice spent a greater proportion of their time in the quadrant that had previously contained the food-containing cup resulting in a discrimination index of 0.64±0.1 compared to the quadrant that previously contained the empty cup (Figure 2A, *Test*, hatched yellow bars; p<0.01). These data suggest that the mice had learned to associate a specific location with food and that an engram representing the memory of this association had been established. We also monitored the length of time interval (Figure S2B, red lines) for this memory to disappear. We found that the animals no longer spent a greater period of time in the quadrant where food had been located if the Test was performed 72 hours after the training, or longer. These data show that the hippocampus can establish an engram linking spatial location to the presence of food. The data to this point also showed that the state of activation of neurons in the hippocampus is altered by changes in nutrition and that glutamatergic neurons in hippocampus can regulate food intake. Because, there are a number of glutamatergic subtypes in hippocampus, we next set out to identify a specific hippocampal subpopulation that could recapitulate these findings.

### Identification of a Food-Sensitive Hippocampal Subpopulation

We set out to identify molecular markers for neurons in the hippocampus whose activity had been altered by the conditions that led to the establishment of this engram using PhosphoTrap, a method that enables profiling cells based on a change in their state of activation (Knight et al., 2012). We began by performing immunohistochemistry for c-fos and pS6 in the control (*Context*) and experimental groups (*Context+Food*) as well as in naive, fasted mice that had not been placed in the chamber with the food cups (Figure 2B-D and Figure S3A; Knight et al., 2012). When compared to naive and Context, the Context+Food group showed increased numbers of hippocampal pS6-positive cells in CA3 and the hilar region (Figure 2B-D and Figure S3A). As previously shown, overlap between c-fos and pS6 activation enables the use of anti-pS6 to precipitate polysomes from activated neurons in order to identify molecular markers for that population (Knight et al., 2012). The availability of molecular markers enables functional studies of the role of the specific neural subpopulations that are identified. The method makes use of the fact that the level of phosphorylation of the S6 ribosomal protein is highly correlated with the state of activation of neurons. Thus, after an acute or chronic stimulus RNAs that are enriched after precipitation of polysomes with a phospho-specific antibody to S6, mark cells that were activated (Knight et al., 2012). Similarly, RNAs that are depleted after polysome precipitation mark cells that have been inhibited. Homogenates of dorsal hippocampi were prepared from *Context* and *Context+Food* mice, immunoprecipitated (IP) with a phosphoS6-specific antibody and the precipitated polysomal RNA was sequenced. The enrichment of each gene was calculated as the number of reads in the precipitated RNA relative to the total (IP/INP; Figure 2E). As expected, we found enrichment for activity-related genes (arc, fosB and cfos; Figure S3B) and a depletion of glial markers (Figure 2F; gfap, aif1 and mal) in both samples.

We next analyzed the RNAseq data to identify genes that were significantly enriched ~2 fold or greater in the precipitated polysomes from hippocampal homogenates prepared from the Context+Food group vs. the Context group (q value <0.05; Figure 2G and Table S1 and S2). We found enrichment for several markers for hilar neurons in the hippocampus including Drd2 (~4.5 average fold; Figure S3C, Figure 2G and 2H). Drd2 was more enriched in the Context+Food experimental group relative to animals in the Context group and also compared to naïve animals (Figure 2I; 4.5 average fold in Context+Food vs. 1.5 average fold in Context vs 1.7 average fold in naive, q<0.05). These results were confirmed using Taqman analysis of Drd2 RNA in the pS6 IP samples from Context+Food animals (~4fold; Figure S3D).

The activation of Drd2*+* neurons in the Context+Food group was then confirmed using dual immunohistochemistry for pS6 expression in a Drd2-cre mouse injected in the hilar region with a virus carrying a cre-dependent mCherry reporter (Drd2-mCherry). Co-localization of Drd2-mCherry and pS6 in the hippocampus was significantly increased in the *Context+Food* group compared to the *Context* group (Figure 2J) while differential expression of pS6 in *Context+Food* vs *Context* was not seen in mCherry-negative cells (Figure 2J; non-Drd2). This suggests that while there are other hippocampal cells that can respond to the context and/or food, the Drd2 neurons respond significantly more to the presence of food. Because Drd2+ neurons (D2R) are active in response to food, we hypothesized that these neurons might play a role to regulate food intake.

Prior reports have shown that D2R neurons in the hippocampus are glutamatergic and thus might represent a subset of the CamkIIa+ neurons we showed to regulate food intake (Etter and Krezel, 2014). We confirmed this by co-injecting an AAV5-Ef1a-DIO-YFP and AAV5-CamkIIa-mCherry in the dorsal hippocampus of Drd2-cre mice and analyzing the co-localization of Drd2-YFP and mCherry (CamkIIa) in the injected area (Figure S4A). Our previous data (see Figure 2I) showed that Drd2 mRNA is significantly more enriched in mice exposed to food compared to mice exposed only to a novel environment. Thus, we first assayed the levels of c-fos expression in D2R neurons in fed and fasted animals (Figure S4B) and found that c-fos expression in D2R neurons is reduced in fasted relative to fed mice. Since overnight fasting was shown to alter water intake and cortisol levels in mice, we also analyzed c-fos expression in D2R neurons in response to water deprivation and a single corticosterone injection (530 nM; Figure S4C; Jensen, 2013) but no significant effect was observed. We also observed in our studies that D2R neurons population in the hilar region is composed of hilar mossy cells and CA3c cells (Scharfman, 1991) and both cell types’ activities are modulated by nutritional changes.

As mentioned, in the comparisons between the *Context* (control) and *Context+Food* (experimental) the animals had been placed in the chamber with food for only five minutes (Figure 2A). Because the limited food intake during this brief interval is not sufficient to induce satiety in fasted mice, we considered the possibility that D2R neurons might be activated by a sensory cue associated with food rather than by caloric intake. We thus tested whether these neurons might be regulated by visual or olfactory stimuli coming from the food in the cups. To test this, we placed fasted mice into a chamber to which they had been habituated with either an empty, clean glass jar (control) or an empty glass jar that had been previously filled with chow (smell). However, once the chow is removed, there is a residual odor allowing us to assess the role of olfactory cues. To test for visual cues, we enclosed food pellets in a tightly sealed glass jar (to block release of odorants) (Figure S4D). Mice are able to discern familiar shapes and visual cues (Huberman and Niell, 2011), and thus should be able to respond to the (familiar) shape of pellets from their home chow. We then quantitated c-fos immunoreactivity in D2Rpositive cells in response to these olfactory and visual stimuli (Drd2-YFP; Figure S4D). While the presence of an empty, cleaned jar failed to induce c-fos expression in D2R neurons (Figure S4D, middle panel, control), the sight of food significantly induced c-fos expression in D2R neurons (Figure S4D; middle panel, sight). Although not significant, a positive trend was observed in mice exposed to the smell of food (Figure S4D, middle panel, smell). These data suggest that D2R neurons can be activated by visual cues and possibly olfactory cues associated with the presence of food. These data prompted us to evaluate whether these neurons could modulate food intake and possibly enable animals to associate sensory cues with food location.

### Functional Analysis of hippocampal D2R neurons

We tested the function of D2R neurons by bilaterally injecting AAVs containing either the cre-dependent activating (hM3Dq; Figure 3, blue bars) or inhibitory (hM4Di; red bars) DREADDs into the hilar region of the hippocampus. Food intake was measured acutely (Figure 3B, 24h, Acute panels) or chronically (8 days; Chronic panels) after an i.p. injection of either CNO or saline. D2R neurons have a baseline firing rate of 1-2Hz and in control studies we found that application of CNO in D2R neurons expressing hM3Dq induced neuronal depolarization and increased their firing rate to 2-8Hz while D2R neurons expressing hM4Di were hyperpolarized by CNO (Figure S5A-D). Mice receiving the inhibitory hM4Di, showed decreased c-fos expression in D2R neurons in the hippocampus after CNO injection (Figure S5E and S5F) while CNO increased c-fos in mice in D2R neurons transduced with the activating hM3Dq DREADD (Figure S5G). We next tested the effect of these treatments on food intake.

**Figure 3.**
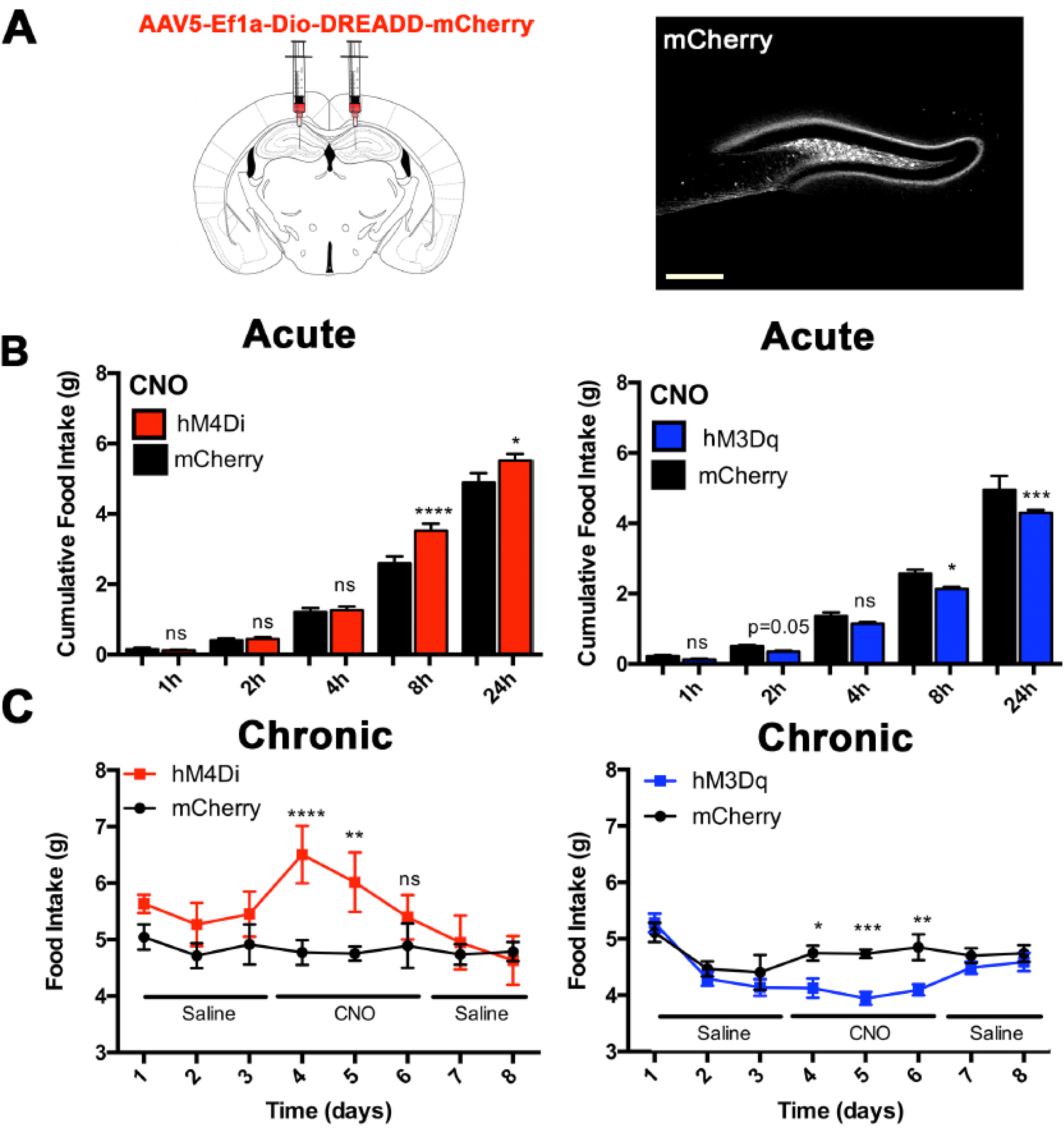
Analysis of the role of Drd2 neurons in food intake. (A) Schematic and representative image of AAV5-Ef1a-DIO-DREADD (hM3Dq and hM4Di)-mCherry expression in the dorsal hippocampus of Drd2-cre mice. Scale bars, 200 *μ*m. (B) Left panel, acute cumulative food intake (g) after a single injection of CNO (i.p; 1 mg/kg) in hM4Di-injected mice (red bars; n=11) and mCherry-injected mice (black bars; n=15). One-way ANOVA with Bonferroni correction, *p<0.05 and ****p<0.0001. Acute (right panel) cumulative food intake (g) after a single injection of CNO (i.p; 1 mg/kg) in hM3Dq-injected mice (blue bars; n=12) and mCherryinjected mice (black bars; n=20). One-way ANOVA with Bonferroni correction, *p<0.05 and ***p<0.001. (C) Left panel, chronic food intake (g) after 3 days of saline injection, 3 days of CNO injection (i.p; 1mg/kg), 2 days of saline injection in hM4Di-injected mice (red lines; n=11) and mCherry-injected mice (black lines; n=11). One-way ANOVA with Bonferroni correction, ****p<0.0001 and **p<0.01. Right panel, chronic food intake (g) after 3 days of saline injection, 3 days of 12/12hr CNO injection (i.p; 1mg/kg), 2 days of saline injection in hM3Dq-injected mice (blue lines; n=20) and mCherry-injected mice (black lines; n=12). One-way ANOVA with Bonferroni correction, ***p<0.001, **p<0.01 and *p<0.05.

Chemogenetic inhibition of D2R neurons acutely increased food intake (Figure 3B, red bars, p<0.05) after 24 hours and chronic inhibition of these neurons also increased food intake (Figure 3C; left panel, p<0.0001 in day 1 and p<0.01 in day 2). In contrast, chemogenetic activation of D2R neurons decreased food intake both acutely (Figure 3B, right panel, p<0.001), and chronically (Figure 3C, right panel; p<0.05 in day 1, p<0.001 in day 2 and p<0.01 in day 3). Saline injection to DREADD-expressing Drd2-cre mice or CNO injection in mCherry-expressing Drd2-cre controls did not significantly alter food intake (Fig S5H and S5I and Figure 3B and 3C, black bars and symbols, respectively). Optogenetic modulation of D2R neuronal activity in a 20-min feeding session also showed the same effects as seen in the chemogenetic study (Figure S5J-L, p<0.05). These data show that activation of D2R neurons in the hippocampus both respond to sensory appetitive cues and also promote satiety.

### Mapping and Functional Testing of Inputs to D2R neurons

The hippocampus is known to receive inputs from several brain regions that modulate its function (Amaral and Witter, 2002). To identify brain regions that project to D2R neurons, we injected a cre-dependent retrograde tracer, pseudorabies virus (PRV) expressing a tdTomato reporter into the dorsal hippocampus of Drd2-cre mice (PRV-lsl-tdTomato; Figure 4A-D, Pomeranz et al., 2017). This virus propagates retrogradely and serially labels presynaptic neurons (Pomeranz et al., 2017). Using this method, we found that D2R neurons receive inputs from contralateral neurons of the dorsal hippocampus (Figure 4C, in average 28 tdTomato+ cells per hippocampus, n=3) and dense inputs from neurons in the superficial and middle layers of the lateral entorhinal cortex as well as a sparse population of neurons in the perirhinal and ectorhinal cortices (overall 164 tdTomato+ neurons, n=3, Figure 4D). To confirm this, we injected an anterograde tracing herpes simplex virus (Lo and Anderson, 2011) into the LEC of Drd2-YFP mice and found co-localization of YFP and HSV-tdTomato (Figure 4E and 4F; in average 17 tdTomato+/YFP+ neurons per hippocampus or 56.6% of D2R+ neurons, n=3). Finally, we used a cre-dependent rabies virus as a cell-specific monosynaptic tracer to confirm that neurons in the LEC project to Drd2 neurons (Fig. S6A; in average of 14 YFP+ neurons per slice in the LEC, n=4).

**Figure 4.**
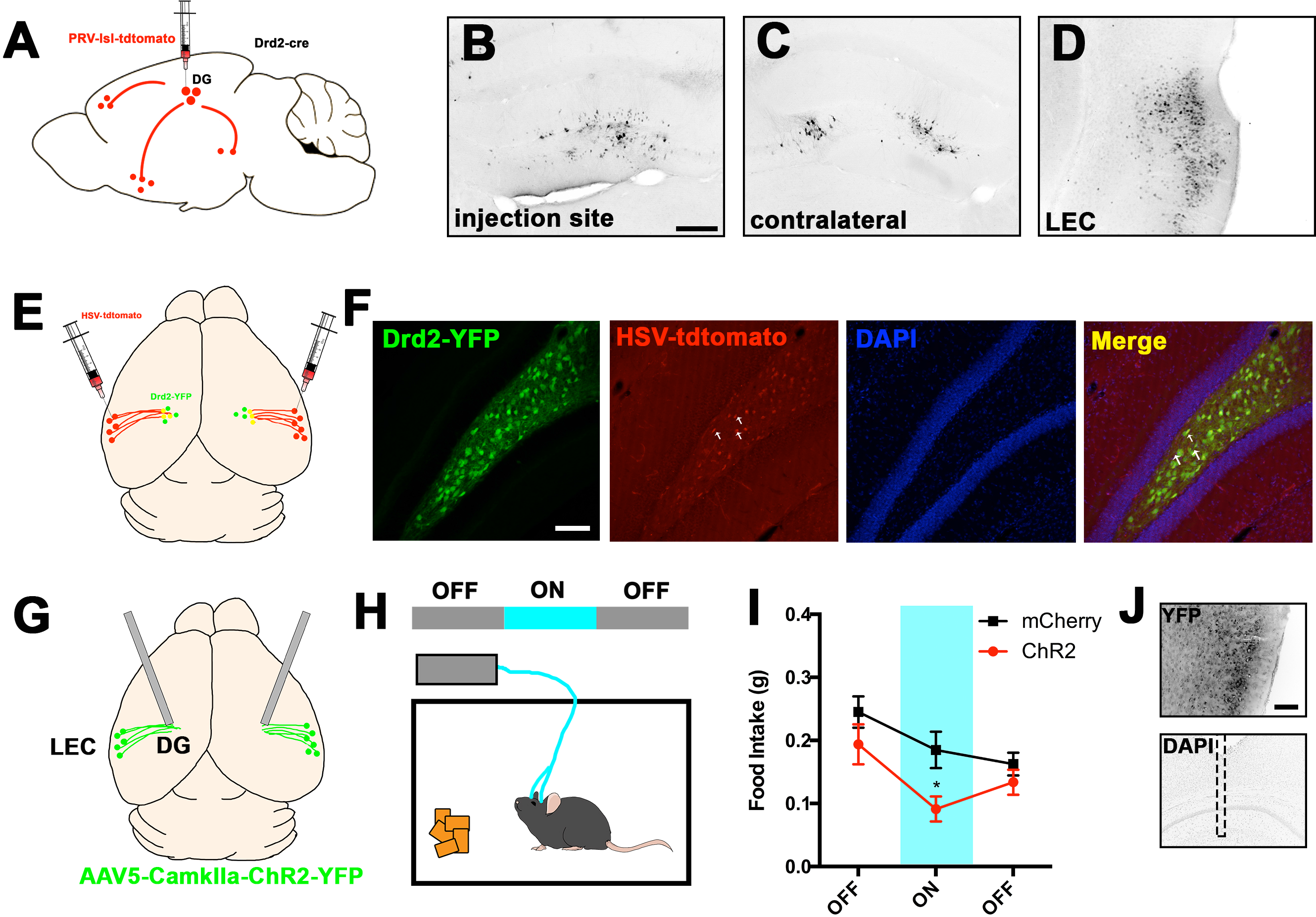
Anatomical mapping of D2R neurons inputs. (A) Schematic drawing of retrograde, transneuronal projection mapping of neuronal inputs to D2R neurons using PRV-lsl-tdTomato in Drd2-cre mice. (B) TdTomato staining of PRV injection site, (C) contralateral side and (D) LEC (lateral entorhinal cortex). Scale bars, 200 *μ*m. (E) Schematic drawing of anterograde projection mapping of neuronal outputs from LEC in Drd2-YFP mice. (F) Images showing staining for Drd2-YFP (green), HSV-tdTomato (red) and nuclei (DAPI; blue). Last right panel show the merged images. White arrows represent YFP+ and tdTomato+ cells. Scale bars, 200 *μ*m. (G) Schematic drawing showing bilateral injection of AAV5-CamkIIa-ChR2-YFP virus in the LEC and optic fiber implantation above the dentate gyrus (DG) of mice. (H) Schematic representation of a 1-hour OFF-ON-OFF feeding session performed in mice. (I) Food Intake during the 1-hour feeding session in ChR2- and mCherry-injected mice. Unpaired t-test, *p<0.05, n=8. (J) Images showing staining for YFP in LEC and optic fiber placement in DG. Scale bars, 200 *μ*m.

To test whether these neurons were able to modulate D2R neurons activity, we injected LEC neurons with an AAV containing activating DREADDs and analyzed c-fos expression in Drd2-YFP neurons in the hilus (Fig. S6D; white arrows; in average 6 c-fos+ neurons per hilus or 20% of D2R neurons). We then tested whether LEC neurons can also modulate feeding behavior by injecting an AAV vector expressing ChR2 under the control of the *CamkIIa* promoter into the LEC of wild-type mice. Optical fibers where then implanted above the DG to assay the effect of activating projections from LEC to hippocampus (Figure 4G-J and Fig. S6B and S6C). The food consumption of control or ChR2-injected mice was tested after local light exposure in a 1-hour OFF-ON-OFF protocol in which food intake was measured after each session (Figure 4H). Optical stimulation of LEC-DG projections decreased food consumption in fasted mice during the ON epochs, but not during the laser OFF epochs or in control mice (Figure 4I; red symbols, p<0.05). These data show that CamkIIa+ LEC neurons can reduce food intake possibly by activating hippocampal D2R neurons.

### Mapping and Functional Testing of D2R neuronal projections

In order to map the outputs of the D2R+ neurons, we injected an AAV expressing a cre-dependent GFP into the hilar region of Drd2-cre mice (Figure 5A) and analyzed YFP expression in brain sections. Images of Drd2-YFP brains revealed dense intrahippocampal projections of D2R+ neurons (primarily coming from a hilar population) to the granule cell region of the dentate gyrus (Figure 5A, first brain panel). We also observed projections of D2R neurons to the lateral septum (LS, Figure 5A, second brain panel) and the Diagonal Band of Broca (DBB, Figure 5A, third brain panel). YFP expression was not seen in the entorhinal area or other cortical regions (Figure 5A), suggesting that the LEC connection to D2R cells is unidirectional. Labeling of D2R neurons also revealed only a very weak projection to the hypothalamus (Figure 5A, fourth brain panel). This finding raised the possibility that the control of food intake by D2R neurons is not conveyed directly through the hypothalamus.

**Figure 5.**
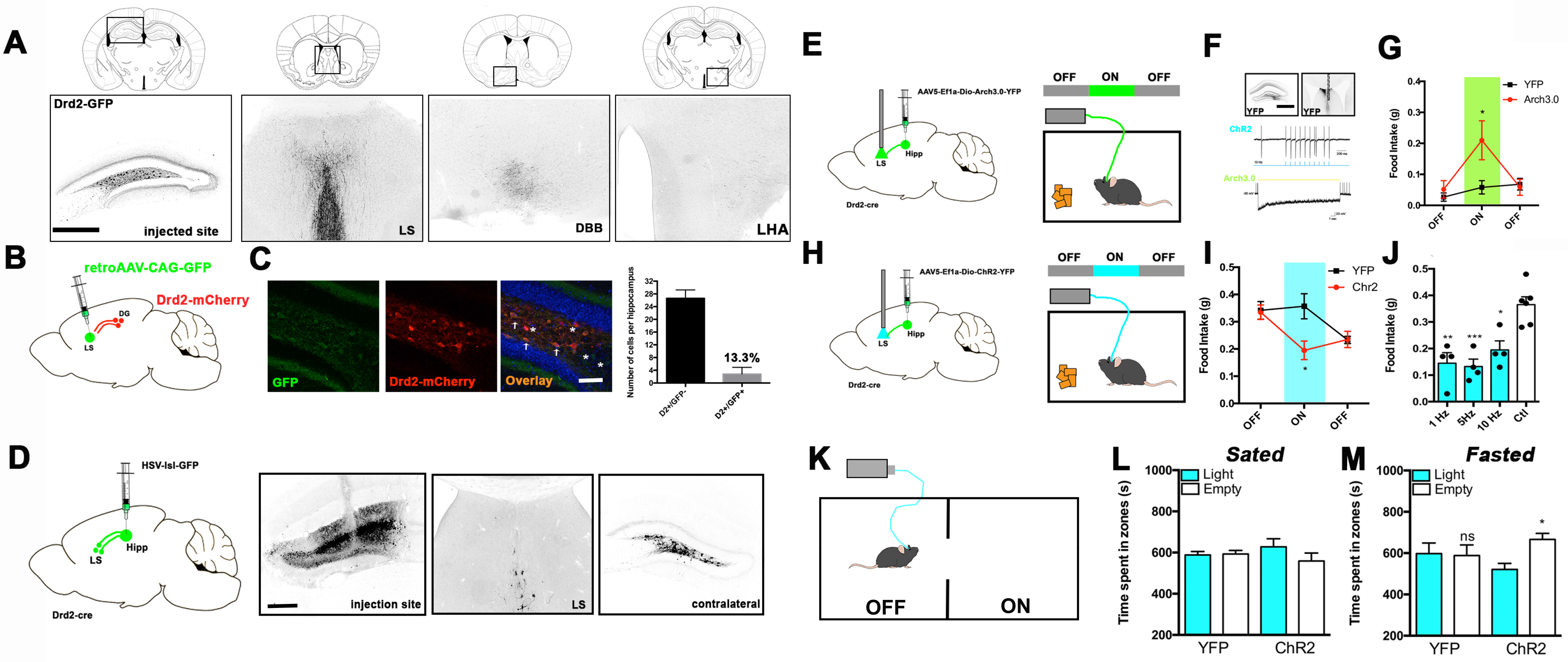
Mapping of D2R neurons downstream projections. (A) Drd2-cre mice were injected with AAV5-Ef1a-DIO-GFP and brain slices were stained with anti-GFP. Brain annotation of axonal projections were done using as reference the Allen Brain Atlas annotated areas: LS (lateral septum), DBB (diagonal band of broca) and LHA (lateral hypothalamic area). (B)Schematic representation of retrograde tracing using retroAAV-CAG-GFP injected in the lateral septal nucleus of Drd2-mCherry mice. (C) Images depicting GFP+ cells that are also Drd2+ in the Drd2-mCherry mouse hippocampus and quantification of Drd2+/GFP+ cells per hippocampus. Scale bars, 200 *μ*m. (D) Schematic representation of anterograde tracing using a cre-dependent HSV-GFP (HSV-lsl-GFP) injected in the dorsal hippocampus of Drd2-cre mice: Hipp (hippocampus) and LS (lateral septum). Right panels show GFP staining in the injection site, LS and contralateral site. Scale bars, 200 *μ*m. (E) The left panel shows schematic drawing showing Drd2-cre mice that were injected in the dorsal hippocampus with AAV5-Ef1a-DioArch3.0-YFP and the right panel shows the schematic drawing of the 1-hour OFF-ON-OFF feeding session. (F) Viral expression and electrophysiological experiment validating the functionality of Arch3.0 and ChR2 expression in Drd2 cells. Scale bars, 400 *μ*m. (G) Measurement of food intake (g) during the 1-hour feeding session. (H) The left panel shows schematic drawing showing Drd2-cre mice that were injected in the dorsal hippocampus with AAV5-Ef1a-Dio-ChR2-YFP and the right panel shows the schematic drawing of the 1-hour OFFON-OFF feeding session. (I) Measurement of food intake (g) during the 1-hour feeding session. Unpaired Student’s t-test, *p<0.05, n=6. (J) Measurement of food intake (g) during the ON epoch of the 1-hour feeding session using different wavelength frequencies (1, 5 and 10 Hz) in ChR2-injected mice and controls. Paired Student’s t-test, *p<0.05, **p<0.01 and ***p<0.001, n=4. (K) Schematic drawing of real time place preference test (RTPP) in Drd2-cre mice containing ChR2-expressing hippocampal cells and optic fibers placed above the LS. (L) RTPP in sated mice or (M) fasted mice. Paired Student’s t-test, *p<0.05, n=8-10.

Analysis of the sites of outputs of the D2R hilar neurons also revealed dense projections to the LS, an area which has previously been shown to control feeding via projections to the lateral hypothalamus (Sweeney and Yang, 2016). We first re-confirmed this projection by injecting a retrograde AAV-CAG-GFP, a retrograde viral tracer (Tervo et al., 2016), into the LS of Drd2-mCherry mice (Figure 5B and 5C). We observed bilateral labeling of D2R cells with GFP 2 weeks post-injection, confirming that D2R neurons project to the lateral septum (Figure 5C, overlay panel, in average 4+ neurons per hippocampus or 13.3% of D2R+ neurons). Finally, we used a cre-dependent herpes virus (HSV-lsl-GFP) as an anterograde polysynaptic transneuronal tracer, to map the circuit from D2R neurons to the LS (Figure 5D; in average 6 GFP+ neurons in the LS per slice), corroborating our previous experiments using immunohistochemistry and thus, defining a synaptic connection between D2R and LS neurons.

The function of the D2R to LS projection was tested by injecting an AAV with cre-dependent expression of either ChR2-YFP or Arch3.0-YFP into the dorsal hippocampus of Drd2-cre mice and placing optical fibers above the LS (Figure 5E-J). Drd2-cre mice injected with a cre-dependent AAV expressing YFP alone were used as controls. Arch3.0 and ChR2 mice were habituated to patch cables and food intake was measured using a 1-hour OFF-ON-OFF feeding session (Figure 5E and 5H, respectively), in which food intake was measured after each 20-minute session. Immunohistochemistry for YFP and electrophysiology confirmed an effect of ChR2 and Arch3.0 expression on D2R neural activity (Figure 5F). We found that in fed mice, inhibiting the D2R projections to the LS increased food intake (Figure 5G, red symbols, p<0.05), while activating this circuit in fasted mice reduced food intake during the ON epochs (Figure 5I, red symbols, p<0.05). No effect was observed during the OFF epochs or in control mice (Figure 5G and 5I, black symbols). These experiments used 10Hz stimulation in ChR2-expressing mice, but a decreased food intake was also observed using lower frequencies (1 and 5Hz; Figure 5J, compare blue bars with white bar). In order to determine whether the satiety effect observed after activation of the D2R-LS circuit was due to changes in motivational valence, we assayed the real-time place preference (RTPP) in ChR2-YFP expressing Drd2-cre mice with optical fibers placed above the LS (Figure 5K-M). We found that after stimulation, mice avoided the light-paired side of the RTPP chamber but only when fasted (Figure 5M, p<0.05). This shows that D2R-LS stimulation is associated with negative valence, but this is only seen when the animals are hungry. Optogenetic activation of this projection failed to alter behavior in an open field test in all experimental groups suggesting that this circuit does not alter anxiety and there were also no changes in total locomotor activity (Figure S7). During and after optogenetic activation of the neurons, the mice failed to show any abnormal behavior and we did not see evidence for seizures, digging and motor or arousal changes (data not shown). These data suggest that D2R neurons can reduce food intake and control motivational valence via projections to the lateral septum.

### The D2R-Lateral Septum Circuit Is Necessary for the Acquisition of an Engram of a Meal

Our findings that the state of activation of D2R neurons is altered by sensory cues associated with food and that these neurons can regulate food intake raised the possibility that these neurons might play a role in the formation of an engram that links food to a specific context (location). To investigate this, we tested whether manipulating the activity of D2R neurons during the acquisition of a memory associating food and spatial location could modulate subsequent behavior (indicative of the retrieval of that memory). In these studies, we employed the same novel task described earlier (see Fig 2A). In this protocol fasted animals are placed in a chamber (to which they had been habituated) with food present in one of the quadrants (Training). When animals receiving food during this training interval, are later placed in the same chamber but without food, they spend significantly more time in the quadrant where the food was previously located (see Figure 2A). We thus asked whether modulation of the activity of D2R neurons during the training period altered the time animals subsequently spent in the quadrant where the food had been previously placed. Alteration of this response would suggest a role for D2R neurons in memory formation (Figure 6).

**Figure 6.**
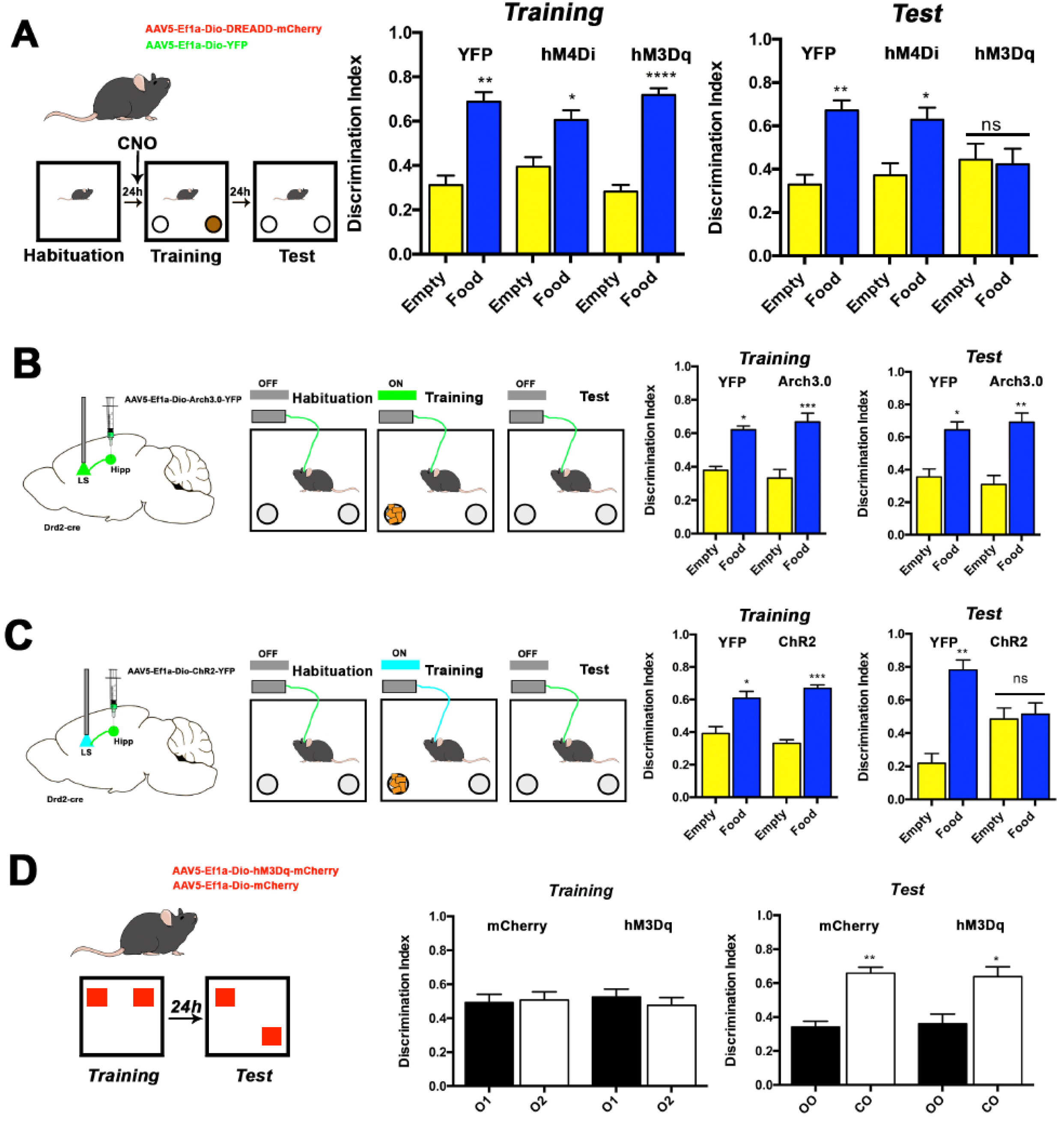
D2R neurons control the acquisition of a memory for the location of food. (A) Left; Schematic representation of a food location task performed in DREADD- or YFP-injected mice. Middle; Discrimination index of YFP-, hM4Di- or hM3Dq-injected mice during Training session. Right; Discrimination index of YFP-, hM4Di- or hM3Dq-injected mice during Test session. Paired Student’s t-test, *p<0.05, **p<0.01 and ****p<0.0001, n=11-15. (B) The left panel shows schematic drawing showing Drd2-cre mice that were injected in the dorsal hippocampus with AAV5-Ef1a-Dio-Arch3.0-YFP and the right panel shows the schematic drawing of the food location task. Middle; Discrimination index of YFP- or Arch3.0-injected mice during Training session. Right; Discrimination index of YFP- or Arch3.0-injected mice during Test session. Paired Student’s t-test, *p<0.05, **p<0.01 and ***p<0.001; n=7-9. (C) The left panel shows schematic drawing showing Drd2-cre mice that were injected in the dorsal hippocampus with AAV5-Ef1a-Dio-ChR2-YFP and the right panel shows the schematic drawing of the food location task. Middle; Discrimination index of YFP- or ChR2-injected mice during Training session. Right; Discrimination index of YFP- or ChR2-injected mice during Test session. Paired Student’s t-test, *p<0.05, **p<0.01 and ***p<0.001, n=8. (D) Schematic drawing of Object Location Task performed in cre-dependent DREADD- or YFP-injected Drd2-cre mice. Middle panel shows the discrimination index during Training and the right panel shows discrimination index during Testing. O1 and O2 depicts similar objects, OO depicts old object and CO depicts changed object. Paired Student’s t-test, *p<0.05 and **p<0.01, n=10.

Consistent with this possibility, we found that chemogenetic activation D2R neurons in hM3Dqexpressing mice reduced the amount of time that trained mice spent in the quadrant that previously contained food (Figure 6; Test, hM3Dq, compare yellow bars with blue bars, no statistical significance and discrimination index of 0.42±0.2 in average). Injection of CNO had no effect on the time spent in this quadrant during the training session when food was present (Figure 6; *Training*) and injection of CNO in control or in hM4Di animals also had no effect on this task (Figure 6, *Test*, YFP and hM4Di). Activating or inhibiting D2R neurons also failed to induce anxiety behavior or alter locomotion as shown using an elevated plus maze (Figure S8A) and an open field test (Figure S8B).

Finally, we tested the role of the projection of D2R neurons from hippocampus to LS on this memory task using optogenetics. An AAV expressing a cre-dependent ChR2or Arch3.0-YFP was injected into the dorsal hippocampus of Drd2-cre mice and optical fibers were placed in the lateral septum (Figure 6B and 6C). We tested the role of this D2R-LS projection by turning the lasers ON during the training session (Training; Figure 6B and 6C) and then subsequently tested whether this affected the time animals spent in the quadrant of the chamber where the food had been previously located. We reasoned that if D2R neurons terminal activity in the LS modulates memory acquisition, a change in the time spent exploring the food cup that previously contained food would be observed in the test session (Test; Figure 6B and 6C). We found that activation of D2R-LS projections reduced the time animals spent in the quadrant of the chamber where the food had been located (Figure 6C, Test, compare ChR2 and YFP, yellow bars and blue bars and discrimination index of 0.48±0.19) while inhibition of this projection had no effect on memory retrieval (Figure 6B, Test panel, compared Arch3.0 and YFP, yellow bars and blue bars, p<0.01 and discrimination index of 0.68±0.16). This suggests that D2R cells play a role in an animals’ ability to associate food with a specific location and furthermore show that D2R projections to the LS can inhibit the acquisition of this memory. In order to assess if D2R neurons role in memory is specific to food, we performed a different non-appetitive, spatial memory task (Object Location Task, Figure 6D). In this experiment, animals are trained in a cage with two similar objects. Then after 24h, the location of one of the objects is changed and the animal’s ability to discriminate between the old and changed object is assessed by monitoring the amount of time the animals spend in the quadrant to which the object had been moved. Notably, activating hippocampal D2R neurons has no effect in the encoding or retrieval of this spatial memory in this task (*Test*, discrimination index of hM3Dq mice was 0.63±0.17, p<0.05). These data show specificity of the circuit we identified for cues associated with food and further suggest that satiety can be accompanied by an inhibition of a circuit that is associated, specifically, with a memory of food location.

## DISCUSSION

Feeding is a complex motivational behavior in which a panoply of sensory inputs are processed to generate an adaptive behavioral response. The successful generation of an adaptive feeding response requires that an animal learn which sensory and interoceptive cues are associated with food availability and its location. Thus, a combination of stimuli that characterize food as safe or unsafe, rewarding or neutral as well as its spatial location is required for the encoding and retrieval of lasting memories of previous experiences of successful foraging (Higgs, 2016; Mela, 2006). However, the identity of neurons in top-down circuits that modulate hunger and satiety through processing of these inputs are still largely unknown, as are the neural pathways through which they act. Because the hippocampus is known to play a key role in episodic memory and learning, we set out to establish the role of specific hippocampal neurons and their neural connections in the acquisition of memories associated with attributes of food including its location, and in the fine-tuning of satiety.

After showing that the activity of hippocampal neurons is sensitive to changes in nutritional state, we used of an unbiased RNA sequencing approach to identify hippocampal neurons that are regulated by food (Knight et al., 2012). We found that neurons in the hippocampus that express the dopamine 2 receptor (D2R) are specifically activated by food and that activation of these neurons suppresses food intake even in mice that had been fasted. We further show that activation of these neurons changes the valence of food and an animal’s ability to associate food, but not an object, with a particular location. Finally, we show that these neurons receive inputs from the LEC and in turn regulate food intake, valence and the acquisition of the memory of food location via projections to the LS.

The hippocampus is subdivided into several molecularly defined areas and is involved in diverse behaviors. Hippocampal cells are necessary for encoding time, location in space and for the encoding and retrieval of episodic memories (Eichenbaum, 2014; Hartley et al., 2013). Thus, a specific ensemble of cells in the hippocampus can be activated by specific experiences (e.g. footshocks received in a specific context), creating an engram, or memory trace of that experience (Ramirez et al., 2013). Indeed, it was shown that an engram from the hippocampus that encodes context modulates the activity of other brain areas, such as the amygdala, to encode the valence of that experience (Ramirez et al., 2013). While previous pharmacological experiments have suggested that the hippocampus can also regulate food intake in mammals (Parent, 2016), neither the identity of the neural populations responsible for this effect have been determined nor had the role of hippocampus in the acquisition of memories associated with food location been established.

We found that most D2R neurons in the hippocampus are localized in the hilar region, with a much sparser D2R+ population in the CA3c layer (Scharfman, 1991). The activity of D2R neurons are also altered by changes in food availability primarily by visual and, to a lesser extent, olfactory characteristics of chow (Figure S4D). We further found that D2R neurons are silenced by fasting, that their activation reduces food intake and finally that this activation is associated with a negative valence as assessed using RTPP (Figure 5K). An analogous set of responses is also observed in CCKexpressing neurons in the nucleus of tract solitarius (CCK^NTS^) that project to the parabrachial nucleus (PBN). CCK^NTS^ neurons are activated following a meal and induce satiety in mice. Additionally, a projection from these neurons to the PBN is aversive (Roman et al., 2017). Satiety may arise from feeling pleasantly (sated) or unpleasantly full, such as after eating too much food. Our data indicated that D2R neurons in the hippocampus contribute to satiety in a manner similar to CCK^NTS^ neurons.

We also found differences in c-fos expression in other hippocampal regions in fed and fasted mice outside the hilar region, including CA3 and DG (Figure 1A). We observed an effect in food intake using chemogenetics to control the activity of glutamatergic *CamkIIa*-expressing cells in the hippocampus though the kinetics and effect size was slightly different than those seen after D2R activation (Figure 1C-E). These data suggest that while D2R neurons represent a functionally important subset of hippocampal cells involved in feeding behavior, it is possible that other cell populations in the hippocampus also play a role to regulate food intake.

Activity mapping using c-fos as a marker for neural activation showed that D2R neurons can also be activated by inputs from glutamatergic *CamkIIa*-expressing neurons in the LEC. While a direct excitatory effect is consistent with the fact that the LEC neurons are glutamatergic, further studies will be necessary to determine whether or not this is a direct effect. The entorhinal cortex is known to convey sensory information from gustatory, olfactory and visual cortices (Li et al., 2017; Scaplen et al., 2017; Seubert et al., 2014) and it too shows changes in its activity in response to changes in metabolic state (Nectow et al., 2017). As mentioned, we found that within the first minutes of exposure to food, D2R neurons are activated by olfactory and visual stimuli raising the possibility that a population of *CamkIIa*-expressing neurons in the LEC might convey relevant sensory cues reflecting the availability of food to the D2R neurons. Further studies will also be required to determine the molecular identity of these putative LEC neurons and whether they convey sensory and/or interoceptive cues to regulate feeding behavior.

The circuit identified herein is also responsive to dopamine as it expresses D2R. This particular receptor is, mostly, a Gi-coupled receptor and thus, dopamine signaling should decrease neuronal activity. However, application of a D2R agonist increases mossy cells excitability (Etter and Krezel, 2014), suggesting that the signaling mechanisms downstream the D2R receptor in hippocampal hilar cells are not completely understood. D2R neurons spontaneously fire in fed animals (Figure S5A-D) and are relatively silent during fasting periods (Figure S4B). In fed mice, D2R neurons are active and our data indicate that they function as part of a circuit that can drive satiety in mice (Figure 3). It is also possible that interoceptive cues as well as other sensory inputs besides those associated with food availability could also regulate the activity of these cells. For example, several hormones have been shown to modulate feeding behavior and the hippocampus expresses the receptors for leptin and ghrelin (Kanoski and Grill, 2015). Furthermore, intrahippocampal injection of leptin has antidepressant effects (Lu et al., 2006) and both ghrelin and leptin have also been shown to modulate memory formation (Kanoski et al., 2013; Kanoski and Grill, 2015) and thus, could play a role in modulating the strength of food-related cues to induce satiety and/or memory formation. Further studies will be necessary to assess whether these and other signals that convey information on the systemic state of energy balance can also influence the activity and function of D2R neurons or the inputs to these neurons from the entorhinal cortex.

We also characterized the functional outputs of D2R neurons and found out that D2R neurons promote satiety by activating cells in the LS (Figure 5). D2R cells directly project to the LS and DBB (Figure 5A), two regions in which there are numerous cholinergic neurons (Herman et al., 2016). Both the LS and the DBB have been implicated in feeding and project directly to several hypothalamic nuclei including the lateral hypothalamic area (Herman et al., 2016; Sweeney and Yang, 2016). Thus, both areas could be inducing satiety after hippocampal D2R stimulation. Determining this will require that the specific populations in the LS and DBB that are activated by D2R neurons be identified.

As mentioned, projections from the LS to neurons in the lateral hypothalamus can influence feeding and neurons in the lateral hypothalamic area (LHA) also receive a set of hormonal and metabolic signals (Domingos et al., 2013a; Leinninger et al., 2011; Li et al., 2015; Sweeney and Yang, 2016). The LHA is known to contain neuronal populations that regulate feeding, motivation, arousal and reward (Berridge et al., 2010; Domingos et al., 2013b), and in turn these LHA neurons send projections to brain areas associated with reward and also cognitive processing, such as the hippocampus (Lima et al., 2013; Stuber and Wise, 2016). It is thus possible that there are reciprocal, indirect, interactions from the hippocampus to the lateral hypothalamus directly or via the lateral septum.

The ability to encode a robust memory denoting the location of food has important adaptive value because a stronger memory would increase the chances of successful foraging and minimize the loss of energy associated with inefficient foraging. The salience of an experience increases the probability of memory formation and, hunger is an especially salient stimulus. Here we show that the exposure to sensory stimuli associated with food, creates an engram in the hippocampus of which D2R neurons are a cellular component. We find that D2R neurons regulate the acquisition of a food-specific memory that is read out using a highly robust behavior, as assayed by the novel Food Location Test we developed, possibly by interacting with other hippocampal cells that encode spatial cues. D2R cells also can encode place fields and thus might also contribute for food-place associations (Senzai and Buzsáki, 2016). We believe D2R cells are able to respond and regulate the encoding of the location of rewarding stimuli, contributing to navigation in goal-directed tasks (Danielson et al., 2017). D2R cells also regulate valence through a LS pathway in fasted, but not sated mice (Figure 5M). This suggests that internal states also contribute, at some level, to how cells in the D2R-LS circuit, or cells that are downstream of it, sense and process reward (Cassidy and Tong, 2017). This might contribute to satiety itself and memory encoding by linking food to an unpleasant sensation (Cassidy and Tong, 2017). This change in valence might also explain the memory deficts observed in our Food Location task after D2R activation. The regulation of valence might also occur via a feedback loop that relies on the projections of D2R neurons to LS (Figure 6) or even through the interaction of LS cells to other areas that project to the hippocampus and control reward, such as the amygdala (Holland, 2004; Zheng et al., 2017). However, the precise mechanisms that couple valence, sensory and cognitive processing to behavior are as yet unknown. Thus, a fuller understanding of the neural mechanisms by which high-order brain areas control these processes represent an important area for further inquiry. The results of these experiments provide an important starting point for this line of investigation.

In summary, we show that D2R neurons are an important cellular component of an LEC->D2R->LS circuit that senses nutritional state and regulates satiety. Our data further indicate that D2R neurons play a role in satiety and appetitive memory processing by controlling valence, and thus, conferring feelings of unpleasantness to food. These studies could have implications for understanding how sensory cues, mental imagery and plainly, thinking about food can influence hunger in mammals.

## AUTHOR CONTRIBUTIONS

E.P.A. and J.M.F conceived and designed the study. E.P.A. performed and analyzed molecular profiling, bioinformatics, feeding and behavioral experiments. J.C performed slice electrophysiology studies. L.P. performed anterograde and retrograde tracing studies using PRV and HSV. M.S helped with feeding experiments using chemogenetics and optogenetics. S.S. helped with PhosphoTrap assay and discussions. K.D provided technical assistance and P.G. assisted with data analysis and discussion. E.P.A and J.M.F wrote the manuscript with input from all authors.

## ACKNOWLEDGMENTS

This work was supported by the JPB Foundation (J.M.F.), and the Human Frontiers in Science Program (E.P.A.). E.P.A is supported by the Long-Term Human Frontiers in Science Program Fellowship and the Kavli NSI Young Investigator Grant. We would like to thank Dr. Eric Nestler for the kind gift of Drd2cre mice. We also wish to thank The Rockefeller University Genomics and Bio-Imaging Resource Centers.

## STAR*METHODS

Detailed methods are provided and include the following:

- KEY RESOURCES TABLE
- CONTACT FOR REAGENT AND RESOURCE SHARING
- EXPERIMENTAL MODEL AND SUBJECT DETAILS
- METHOD DETAILS

- Stereotaxic injections and Optic Fiber Implantation
- Feeding experiments using chemogenetics and optogenetics
- Immunohistochemistry, image processing and cell counting
- Food Location Task
- Object Location Task
- PhosphoTrap
- Water deprivation and Corticosterone Injection
- RNA sequencing and qPCR analysis
- Real-Time Place Preference
- Sensorial Task
- Retrograde and Anterograde Tracing
- Locomotion assessment for optogenetics and chemogenetics
- Elevated Plus Maze for chemogenetics
- Slice electrophysiology
- QUANTIFICATION AND STATISTICAL ANALYSIS
- DATA AND SOFTWARE AVAILABILITY

- Data Resources

## CONTACT FOR REAGENT AND RESOURCE SHARING

Further information and requests for reagents may be directed to, and will be fulfilled by the corresponding author Jeffrey Friedman.

## EXPERIMENTAL MODEL AND SUBJECT DETAILS

Mice were housed according to the guidelines of the Rockefeller’s University Animal Facility, in a 12 hr light/dark cycle and with free access to standard chow diet and water, in exception when specified in the text. C57BL/6J background males and females at age 8-12 weeks were used throughout this study and all animals experiments were approved by the Rockefeller University IACUC following the National Institutes of Health guidelines for the Care and Use of Laboratory Animals. For behavioral studies, only male mice were used. C57BL/6J wild-types were purchased from Jackson Laboratories (Bar Harbor, ME). Drd2-cre mice were a gift from Dr. Eric Nestler, from Mount Sinai School of Medicine.

## METHODS DETAILS

### Stereotaxic injections and Optic Fiber Implantation

Mice were anesthetized with 2% isoflurane, placed in a stereotaxic frame (Kopft Instruments). Eye ointment was applied to the eyes and a subcutaneous injection of meloxicam (2 mg/kg) was given to each mouse after surgery for up to 2 days. Hair was shaved, the scalp was disinfected with iodine solution and an incision was made. The craniotomy was performed using a dental drill (Dremel). A hamilton syringe (Cat# 84851; 85RN; 26s/2”/2) was used to infuse the virus at a rate of 0.1 μL/min at a total volume of 0.5 μL. After viral infusion, the needle was kept at the injection site for 10 min and then slowly withdrawn. High titer (10^11^) viral preparations were purchased from University of North Carolina at Chapel Hill Vector Core and injected unilaterally at the following coordinates: for lateral septum: from bregma, AP: +0.86 mm, ML:0.0 mm, DV:−3.00 mm. And bilaterally: dorsal hippocampus: from bregma, AP: − 1.94 mm; ML: −/+ 1.25 mm, DV: −2.04 mm with a 6° degree angle; for lateral entorhinal cortex injections: from bregma, AP: −2.8 mm, ML: −/+ 4.4 mm, DV: −4.3 mm. For all surgeries, mice were monitored for 72h to ensure full recovery and 2-4 weeks later mice were used in experiments. Optic fiber implants (Thor Labs) for optogenetics experiments were implanted to the same coordinates as the viral injections with a −0.5 mm difference. Implants were inserted slowly and secured to the mouse skull using two layers of Metabond (Parkell Inc) followed by a layer of dental cement. Mice were single housed and monitored in the first weeks following optic implantation.

### Feeding experiments using chemogenetics and optogenetics

For acute feeding, control or DREADD-injected mice were single housed and injected with 0.9% saline or clozapine-N-oxide (CNO; Sigma) at 1 mg/kg was intraperitoneally (i.p) before dark phase and cumulative food intake was measured in selected time periods. For chronic feeding, mice were single or double i.p. injected (every 12h) with CNO or saline before dark phase and food intake was measured. At the completion of experiments, control and DREADD-injected mice were sacrificed in order to confirm efficient viral expression. For experiments that require fasting, mice were single housed, and food was removed before dark phase (16-23h of fasting in total). After, fasted mice were perfused transcardially and brains were processed for immunohistochemistry. Control mice (Fed) were sacrificed and processed at the same time as fasted mice. Feeding experiments using mice expressing a cre dependent ChR2- or Arch3.0-YFP were performed in a specific behavior room. Mice were acclimated to the behavior room for a week before experimental assays in a 12/12h light cycle and handled for 3 min daily for 4 days. Implanted optic fibers were attached to a patch cable using ceramic sleeves (Thorlabs) and connected to 473 or 532 nm laser (OEM Lasers/OptoEngine). Laser output was verified at start of each experiment. Blue light was generated by a 473 nm laser diode (OEM Lasers/OptoEngine) and yellow light by a 532 nm laser diode at 5-10 mW of power. Light pulse (10 ms) and frequency (1-10Hz) was controlled using a waveform generator (Keysight) to activate Drd2+ cell bodies, *Drd2+* terminals in lateral septum or *CamkIIa+* terminals in dorsal hippocampus. Animals were sacrificed to confirm viral expression and fiber placement using immunohistochemistry. Each feeding session lasted 80 min and it was divided in 1 trial of 20 min to allow the animal to acclimate to the cage and 3 trials of 20 min each (1h feeding session). During each feeding session, light was off during the first 20 min, on for 20 min and off again for the remaining 20 min. Consumed food was recorded manually before and after each session. To facilitate measurement, 3 whole pellets were added to cups and food crumbs were not recorded. Control mice expressing a cre-dependent YFP or mCherry received the same treatment as ChR2- and Arch3.0-YFP mice. Feeding bouts were recorded using a camera and the Ethovision 9.0 software (Noldus).

### Immunohistochemistry, image processing and cell counting

Mice were transcardially perfused and brains were postfixed for 24h in 10% formalin. Brain slices at 50 μm were taken using a vibratome (Leica), blocked for 1h with 0.3% Triton X-100, 3% bovine serum albumin (BSA), and 2% normal goat serum (NGS) and incubated in primary antibodies (see Key Resources Table) for 24h at 4°C. Then, free-floating slices were washed three times for 10 min in 0.1% Triton X-100 in PBS (PBS-T), incubated for 1h at room temperature with secondary antibodies, washed in PBS-T and mounted in Vectamount with DAPI (Southern Biotech). Antibodies used here were: antipS6 (1:1000), anti-c-fos (1:500), anti-GFP (1:1000), anti-mCherry (1:1000), goat anti-rabbit (Alexa 488 or Alexa 594, 1:1000), goat anti-chicken Alexa488 or Alexa594 (1:1000). For double staining for pS6 and cfos, tissues were serially stained as described previously (Knight et al., 2012). Images were taken using Axiovert 200 microscope (20x objective; Zeiss) or a lnverted LSM 780 laser scanning confocal microscope (20x objective; Zeiss) and c-fos staining were manually quantified using the CellCounter plugin and ImageJ software (NIH). To count *CamkIIa+* and Drd2+ cells, 3 mice were injected with a mix of AAV5-CamkIIa-mCherry and AAV5-EF1a-Dio-YFP. After 2 weeks, mice were sacrificed, and brains processed for immunohistochemistry. *CamkIIa+* cells (red) and Drd2+ cells (green) were counted manually using the CellCounter plugin and ImageJ software (NIH).

### Food Location Task

For the food location task, animals were acclimated to the behavior room were the task was held and handled for 3 min each day for at least 4 days before the task. Mice were habituated for 5 min in a 28 × 28 cm open field arena previously cleaned with 70% ethanol and divided into 4 equal quadrants. After exploring the arena, mice were returned to their home cage and fasted for 16-23h. The next day, mice were returned to the same open field arena containing two clean, empty food cups and allowed to explore the context for 5 min (termed Context). For Context+Food or Training groups, to one of the paper food cups it was added 3-4 standard chow pellets and mice were allowed to explore the *Context+Food* environment for 5 min. For assessing memory consolidation of the location of the food pellets (termed Test), mice were habituated and trained as described above and then returned after training to their home cage, fed and before dark phase started, mice were fasted for an additional 1623h. The next day (Test), mice were allowed to explore for 5 min an open field arena containing two empty paper food cups. Boxes and food cups were cleaned with 70% ethanol in between trials to remove any food crumbles or mice smell. For the food location task using chemogenetic manipulation mice were habituated as described above and 1h before training, CNO (1 mg/kg) was i.p injected. For food location task using optogenetic manipulation, mice were attached to a patch cable through ceramic sleeves and habituated to the arena for 5 min. After, mice were removed, and food pellets were added to one food cup. Mice were then allowed to explore the arena for 5 min under constant (Arch3.0) or pulsed (ChR2) laser stimulation (parameters described in ‘Feeding experiments using chemogenetics and optogenetics’). To analyzed forgetting, time spent in the food quadrant was analyzed periodically at 24h, 72h and 1 week after training. Fasted mice were overnight fasted before training and before each testing session. In all conditions, total distance and time spent in each quadrant was recorded using a camera and the Ethovision 9.0 software (Noldus) and analyzed previously using GraphPad Prism 5.0 (GraphPad). For training and test condition a discrimination index was calculated as the ratio between the time spent in the given quadrant and the sum of time spent at the two quadrants containing cups. Discrimination indexes were not plotted for empty quadrants, as time spent in those were negligible in all experiments. To analyze, c-fos and pS6 expression groups of 4 mice each were sacrificed 1h after each session and immunohistochemistry was performed.

### Object Location Task

For the object location task, animals were acclimated to the behavior room where the task was held and handled for 3 min each day for at least 4 days before the task. Mice were habituated for 5 min in a 28 × 28 cm open field arena previously cleaned with 70% ethanol. During the training phase, two identical objects are place in opposite sides of the arena and mice are free to explore the arena and objects for 10 min. After a 24h delay, mice perform the testing session of the task, in which one of the object’s location is changed and the mice are free to explore the arena for 10 min. For training and test condition a discrimination index was calculated as the ratio between the time spent in the given quadrant and the sum of time spent at the two quadrants containing objects. Discrimination indexes were not plotted for empty quadrants, as time spent in those were insignificant in all experiments. Arena and object were cleaned with 70% alcohol in between subjects and object preference was tested before task.

### PhosphoTrap

Trap was performed according to Knight et al., 2012. Briefly, mice were separated into groups of 4-6 mice per group (termed naive, context, context+food), allowed to perform the food location task and sacrificed 1h after each task. The Context+Food group refers to mice provided with food for 5 min in a chamber they have been habituated to, while Context group refers to animals exposed only to the chamber itself. Naive mice were kept in their home cage until euthanasia. After euthanasia, brains were removed, and hippocampi were dissected on ice and pooled (8-12 hippocampi per experiment). Tissue was homogenized and clarified by centrifugation. Ribosomes were immunoprecipitated by using 4 ug of polyclonal antibodies against pS6 (Invitrogen) previously conjugated to Protein A-coated magnetic beads (Thermofisher). A small amount of tissue RNA was saved before the immunoprecipitation (Input) and both input and immunoprecipitated RNA (IP) were then purified using RNAeasy Mini kit (QIAGEN) and RNA quality was checked using a RNA PicoChip on a bioanalyzer. RIN values >7 were used. Experiments were performed in triplicates for each group. cDNA was amplified using SMARTer Ultralow Input RNA for Illumina Sequencing Kit and sequenced on an Illumina HiSeq2500 platform. As a protocol efficiency control, we also monitor the enrichment for hbb-b1 gene (hemoglobin beta chain, b-1) using qPCR.

### Water deprivation and corticosterone injection

To test the effect of water deprivation, mice were water deprived for 16h but had free access to food. Control mice had access to food and water *ad libitum*. Corticosterone (530 nM) was diluted in saline and i.p. injected in *Drd2*-mCherry mice. Control mice were i.p. injected with saline only. Water deprived mice and controls as well as mice injected with corticosterone or vehicle injection after 1h of injection were transcardially perfused. Brains were processed according to Methods section “*Immunohistochemistry*, *image processing and cell counting*”.

### RNA sequencing and qPCR analysis

RNA sequencing raw data was uploaded and analyzed using BaseSpace apps (TopHat and Cufflinks; Illumina) using an alignment to annotated mRNAs in the mouse genome (UCSC, Mus musculus assembly mm10). The average immunoprecipitated (IP) and Input value of each enriched and depleted genes with a q-value lower than 0.05 were plotted using GraphPad Prism (GraphPad). To validate the RNA Sequencing data, qPCR using predesigned Taqman probes was performed. For qPCR, predesigned Taqman probes (idtDNA) and cDNA was prepared using the QuantiTect Reverse Transcription Kit (Life Technologies). The abundance of these genes in IP and Input RNA was quantified using Taqman Gene Expression Master Mix (Applied Biosystems). Transcript abundance was normalized to beta-actin. Fold of Change were calculated using the ΔΔCt method and results were statistically analyzed using unpaired student’s t-test.

### Real-time Place Preference

Mice were habituated to optic patch cables for 3-5 days as described above and placed on the border between two adjoining homogenous black compartments with white floors and the amount of time spent in each compartment was recorded using video tracking software (Ethovision 9, Noldus). Most mice displayed no preference, but those with >75% side preference on a pre-test were excluded from further study. On the subsequent day, one side was designated light-paired in which active entry triggered photostimulation (10 Hz, 10 ms pulse width, 5-10 mW), using the lasers as described above but controlled by a Mini I-O box from Noldus and a waveform generator (Keysight). Sessions lasted for 20 min and the amount of time spent in each compartment was recorded.

### Sensorial Task

To evaluate c-fos expression in mice exposed to the smell or sight of food, mice were habituated for 4 days to a 28 × 28 cm open field arena and groups of mice were exposed to a cleaned, empty jar (control group), an empty jar previously dipped in standard chow to acquire the smell of food (smell group) or a sealed, cleaned jar containing food pellets inside (sight). All groups had 5 min to explore the arena containing the stimuli above mentioned and 30 min later mice were sacrificed, and brains were processed for immunohistochemistry.

### Retrograde and Anterograde Tracing

PRV-Introvert-GFP is a retrograde tracing virus in which both transsynaptic tracing and GFP expression are initiated only after activation by cre recombinase (Pomeranz et al., 2017). HSV-H129-tdTomato is a mutant of strain HSV-H129 that was produced by passage of HSV-H129-**Δ**TK-TT (Lo and Anderson, 2011) in cre-expressing cells and selection and purification of tdTomato positive plaques (L.E. Pomeranz, unpublished). HSV-H129-**Δ**TK-TG (a kind gift from David Anderson) is a cre-dependent anterograde transneuronal tracer that expresses GFP after cre activation. Viral stocks were aliquoted and stored at −80°C. 500 nl of virus were injected using the coordinates described herein and mice were perfused 3-4 days after. For retroAAV-CAG-GFP (Addgene), mice were perfused 2 weeks after to ensure maximum viral expression. After, brains were processed for immunohistochemistry.

### Locomotion assessment for optogenetics and chemogenetics

For chemogenetics, control or DREADD-injected mice were acclimated to the behavior room for a week and injected with CNO (1mg/kg) 1h before allowed to explore a 28 × 28 cm arena for 15 min. After, mice were returned to its home cage and the arena floor was cleaned in between subjects. For optogenetics, injected mice were attached to a patch cable through a ceramic sleeve and allowed to habituate for in their home cage. After, mice were place in the arena for a 3 × 5 min session consisting in one session of 5 min with laser excitation turned off, 5 min on and 5 min off. During on sessions, laser stimulation was either constant (5-10 mW; Arch3.0) or pulsed (5-10 mW, 10Hz, 10 ms; ChR2). All subjects were recorded using a camera and analyzed using Ethovison 9.0 (Noldus). Locomotion was assessed as total distance plotted using GraphPad Prism 5.0 (GraphPad).

### Elevated Plus Maze for chemogenetics

Control or DREADD-injected mice were acclimated to the behavior room for a week and injected with CNO (1mg/kg) 1h before allowed in the maze. The maze consists in a cross-shaped, elevated from the floor in which two arms are closed with dark walls and two arms are open. Mice were placed at the center of the maze and allowed to explore it for 10 minutes. After, mice were returned to its home cage and the maze floor was cleaned in between subjects. All subjects were recorded using a camera and analyzed using Ethovison 9.0 (Noldus). Time spent in open and closed arms were quantified and plotted using GraphPad Prism 5.0 (GraphPad).

### Slice electrophysiology

Mice aged 9 to 10 weeks old were euthanized with CO2. Mouse brain was then removed and placed in an ice-cold N-Methyl-D-glucamine (NMDG)-containing cutting solution (in mM: 93 NMDG, 2.5 KCl, 1.2 NaH2PO4, 30 NaHCO3, 25 glucose, 20 HEPES, 5 sodium ascorbate, 3 sodium pyruvate, 2 thiourea, 0.5 CaCl2, 10 MgSO4, pH 7.4, 295-305 mOsm). Brain slices (400 μm thickness) containing the dorsal hippocampus were cut by using a VT1000 S Vibratome (Leica Microsystems Inc.). After cutting, slices were allowed to recover in the cutting solution for 15 min at 37 °C and then transferred to the recording solution at room temperature for at least 1 hr before recording. Brain slices were placed in a perfusion chamber attached to the fixed stage of an upright microscope (Olympus) and submerged in continuously flowing oxygenated recording solution containing the following (in mM): 125 NaCl, 25 NaHCO3, 25 glucose, 2.5 KCl, 1.25 NaH2PO4, 2 CaCl2, and 1 MgCl2, pH 7.4, 295-305 mOsm. Neurons were visualized with a 40× water immersion lens and illuminated with near infrared (IR) light. Electrophysiological recordings were performed with a Multiclamp 700B/Digidata1440A system (Molecular Devices). Patch electrodes were filled with the internal solution (in mM: 126 K-gluconate, 10 NaCl, 2 MgSO4, 1 CaCl2, 1 BAPTA, 15 glucose, 19 HEPES, 2 ATP, and 0.3 GTP, pH 7.3, 290 mOsmol). Whole-cell patch-clamp recordings were used to record the spontaneous action potential of mossy cells. Data analysis for the electrophysiological recordings was performed in Clampfit (Molecular Devices).

## QUANTIFICATION AND STATISTICAL ANALYSIS

Sample sizes (n = number of animal per group), statistical tests used, and statistical significance are described in the Figures and the Figure Legends. Unless otherwise indicated, values are reported as mean ± SEM. (error bars). Behavioral and quantification experiments were done in a blinded manner. One-way ANOVA statistical test was followed by post hoc comparisons using Bonferroni multiple comparisons test. Unpaired Student’s t-test was used to compare two groups. Mice with no viral expression were excluded from the final analysis. In figures, asterisks show the obtained statistical significance *p < 0.05, **p < 0.01, ***p < 0.001 and ****p < 0.0001. All statistical analysis was performed using GraphPad Prism 5.0 software (GraphPad).

## DATA AND SOFTWARE AVAILABILITY

### Data Resources

Further requests for data may be directed to and will be fulfilled by the corresponding author Jeffrey Friedman.

## STAR*METHODS

### KEY RESOURCES TABLE

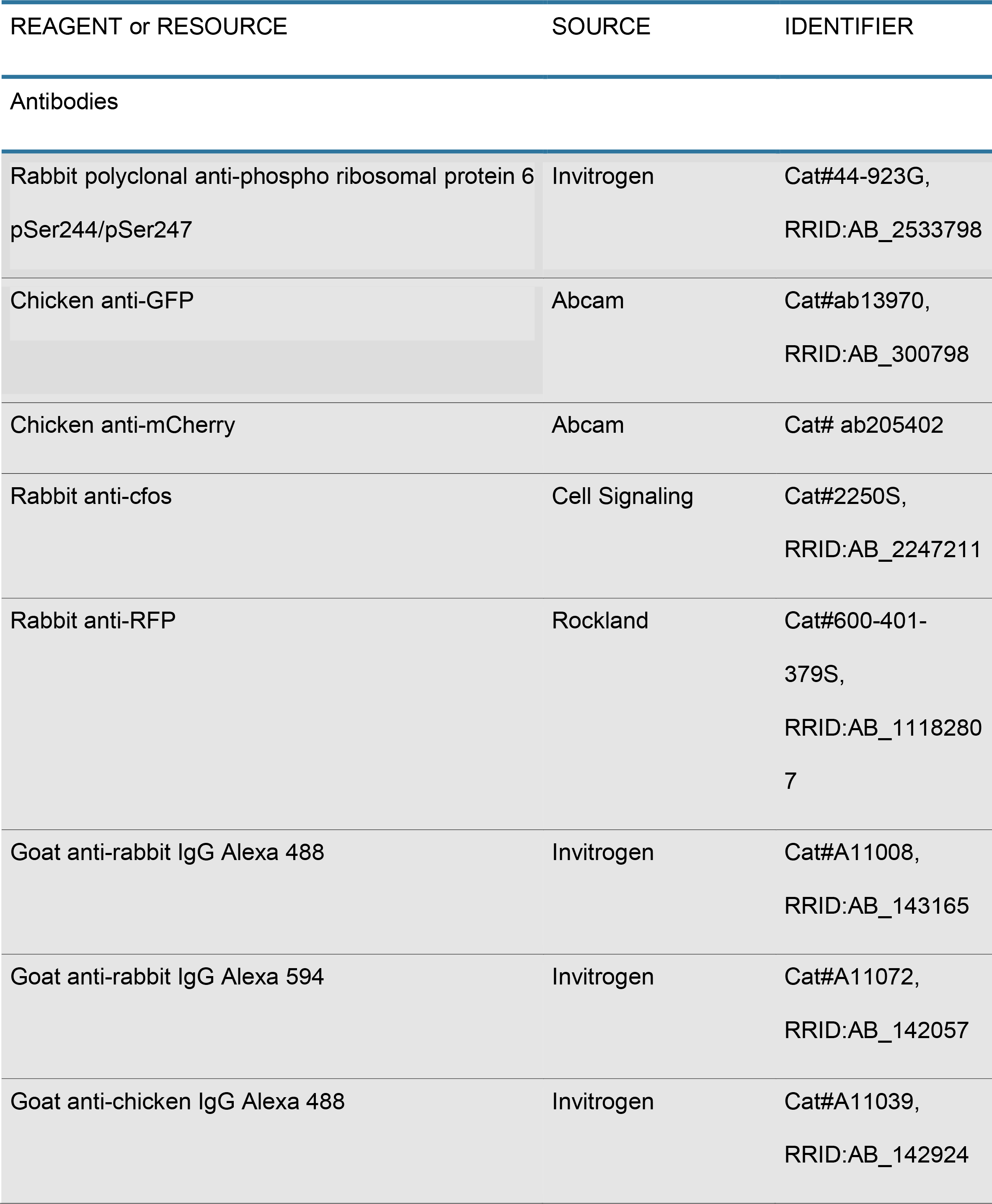

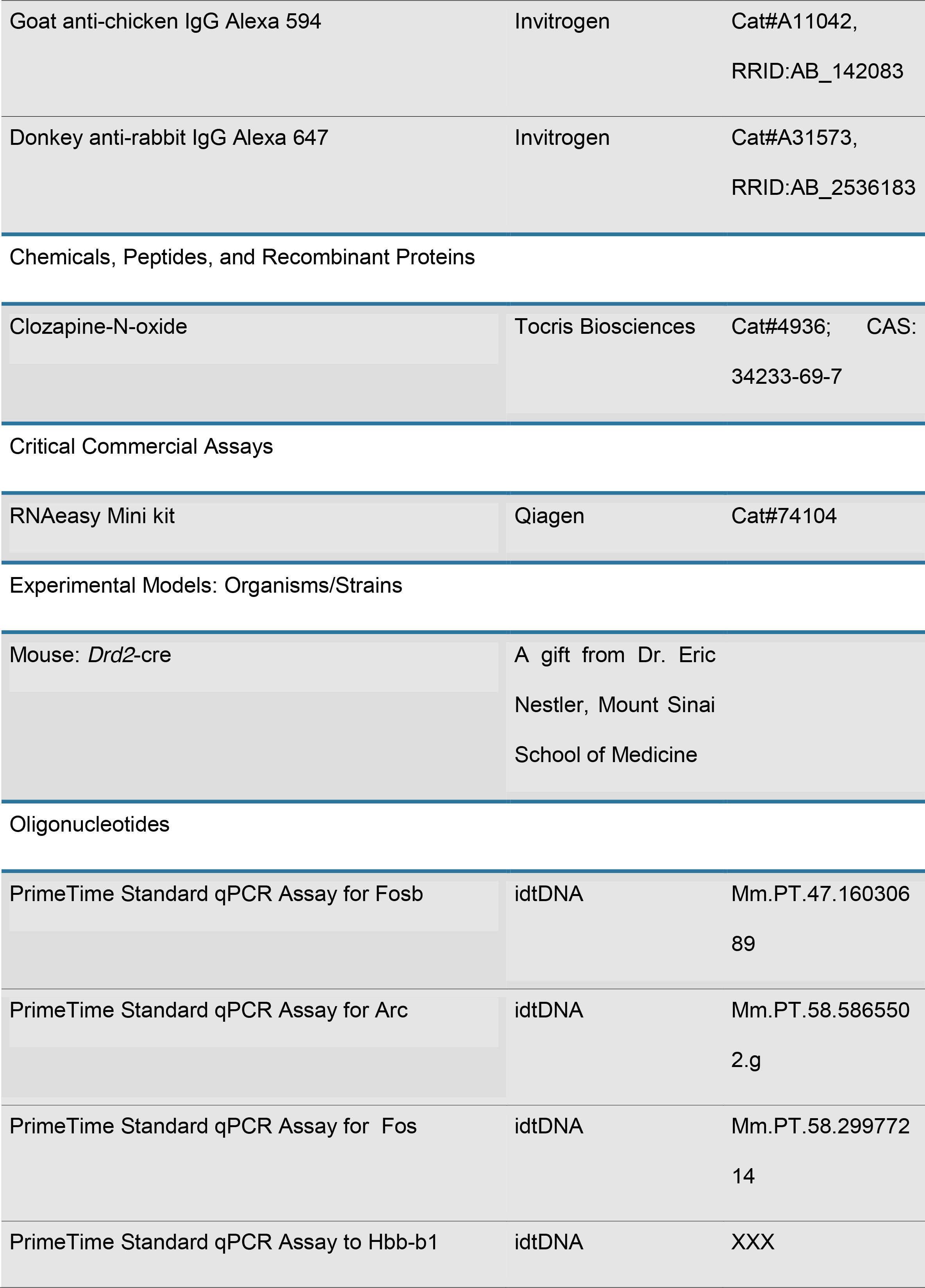

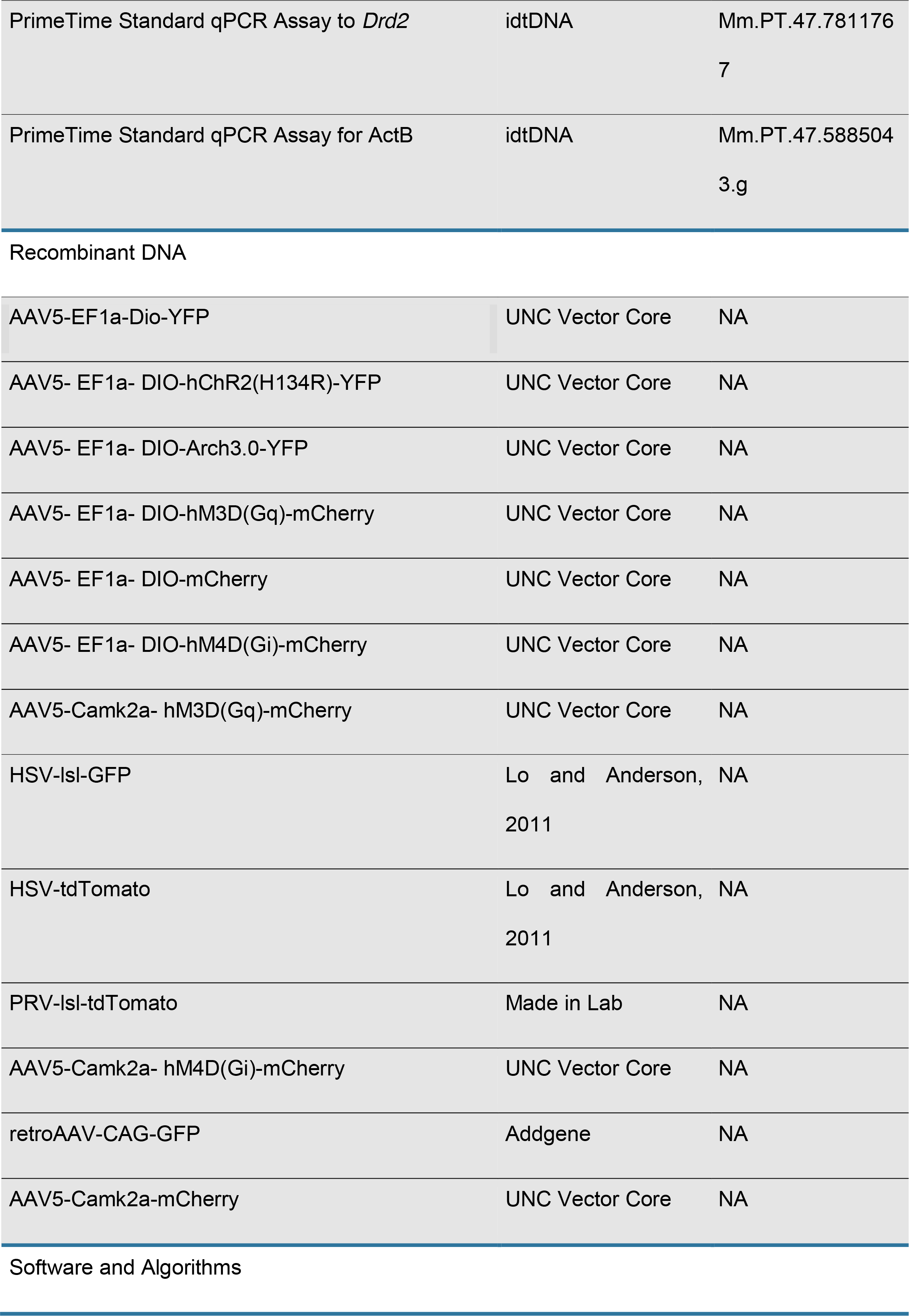

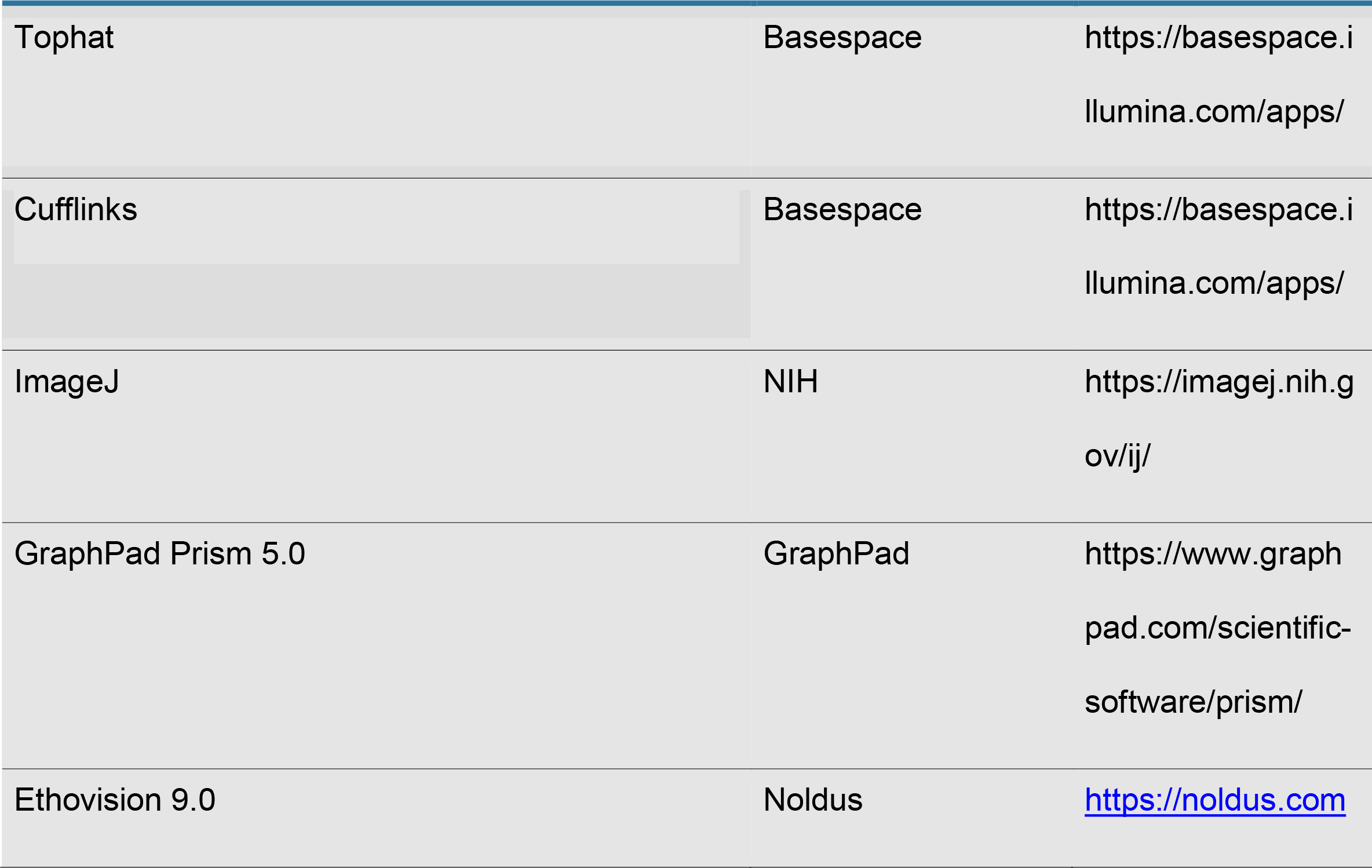

| REAGENT or RESOURCE | SOURCE | IDENTIFIER |
| --- | --- | --- |
| Antibodies |  |  |
| Rabbit polyclonal anti-phospho ribosomal protein 6 pSer244/pSer247 | Invitrogen | Cat#44-923G, RRID:AB_2533798 |
| Chicken anti-GFP | Abcam | Cat#ab13970, RRID:AB_300798 |
| Chicken anti-mCherry | Abcam | Cat# ab205402 |
| Rabbit anti-cfos | Cell Signaling | Cat#2250S, RRID:AB_2247211 |
| Rabbit anti-RFP | Rockland | Cat#600-401-379S, RRID:AB_11182807 |
| Goat anti-rabbit IgG Alexa 488 | Invitrogen | Cat#A11008, RRID:AB_143165 |
| Goat anti-rabbit IgG Alexa 594 | Invitrogen | Cat#A11072, RRID:AB_142057 |
| Goat anti-chicken IgG Alexa 488 | Invitrogen | Cat#A11039, RRID:AB_142924 |
| Goat anti-chicken IgG Alexa 594 | Invitrogen | Cat#A11042,<br>RRID:AB_142083 |
| Donkey anti-rabbit IgG Alexa 647 | Invitrogen | Cat#A31573,<br>RRID:AB_2536183 |
| Chemicals, Peptides, and Recombinant Proteins |  |  |
| Clozapine-N-oxide | Tocris Biosciences | Cat#4936; CAS:<br>34233-69-7 |
| Critical Commercial Assays |  |  |
| RNAeasy Mini kit | Qiagen | Cat#74104 |

Experimental Models: Organisms/Strains
|  |  |
| --- | --- |
| Mouse: *Drd2*-cre | A gift from Dr. Eric<br>Nestler, Mount Sinai<br>School of Medicine |
| Oligonucleotides |  |  |
| PrimeTime Standard qPCR Assay for Fosb | idtDNA | Mm.PT.47.160306<br>89 |
| PrimeTime Standard qPCR Assay for Arc | idtDNA | Mm.PT.58.586550<br>2.g |
| PrimeTime Standard qPCR Assay for Fos | idtDNA | Mm.PT.58.299772<br>14 |
| PrimeTime Standard qPCR Assay to Hbb-b1 | idtDNA | XXX |
| PrimeTime Standard qPCR Assay to *Drd2* | idtDNA | Mm.PT.47.781176<br>7 |
| PrimeTime Standard qPCR Assay for ActB | idtDNA | Mm.PT.47.588504<br>3.g |
| Recombinant DNA |  |  |
| AAV5-EF1a-Dio-YFP | UNC Vector Core | NA |
| AAV5- EF1a- DIO-hChR2(H134R)-YFP | UNC Vector Core | NA |
| AAV5- EF1a- DIO-Arch3.0-YFP | UNC Vector Core | NA |
| AAV5- EF1a- DIO-hM3D(Gq)-mCherry | UNC Vector Core | NA |
| AAV5- EF1a- DIO-mCherry | UNC Vector Core | NA |
| AAV5- EF1a- DIO-hM4D(Gi)-mCherry | UNC Vector Core | NA |
| AAV5-Camk2a- hM3D(Gq)-mCherry | UNC Vector Core | NA |
| HSV-IsI-GFP | Lo and Anderson, 2011 | NA |
| HSV-tdTomato | Lo and Anderson, 2011 | NA |
| PRV-IsI-tdTomato | Made in Lab | NA |
| AAV5-Camk2a- hM4D(Gi)-mCherry | UNC Vector Core | NA |
| retroAAV-CAG-GFP | Addgene | NA |
| AAV5-Camk2a-mCherry | UNC Vector Core | NA |
| Software and Algorithms |  |  |
| Tophat | Basespace | https://basespace.illumina.com/apps/ |
| Cufflinks | Basespace | https://basespace.illumina.com/apps/ |
| ImageJ | NIH | https://imagej.nih.gov/ij/ |
| GraphPad Prism 5.0 | GraphPad | https://www.graphpad.com/scientific-software/prism/ |
| Ethovision 9.0 | Noldus | https://noldus.com |

## SUPPLEMENTAL INFORMATION

Supplemental information includes eight supplementary figures and two tables.

**Supplementary Figure 1. Immunostaining for c-fos in DREADD-injected mice.** (A) Schematic drawing of the experimental design. (B) mCherry (left) and c-fos (right) staining in AAV5-CamkIIa-hM4Di-mCherry injected mice. (C) mCherry (left) and c-fos (right) staining in AAV5-CamkIIa-hM3Dq-mCherry injected mice. Scale, 100 *μ*m

**Supplementary Figure 2. Food Location Task.** (A) Time spent in quadrant during the Training phase of Food Location Task. Yellow bars: quadrant with empty cups; Blue bars: quadrants with food-containing cups; White bars: empty quadrants. (B) Discrimination index of the Testing phase of Food location task in fast (red symbols and lines) mice in a time course of 1 week. Paired T-Test, *p<0.05, ** p<0.01; n=8

**Supplementary Figure 3. PhosphoTrap analysis and gene markers.** (A) Quantification of pS6+ cells in the hippocampus of naïve (white bars), Context (blue bars) and Context+Food (yellow bars) mice. One-way ANOVA with Bonferroni correction, n=3, **p<0.01 and *p<0.05. (B) mRNA levels of activity-dependent genes. (C) Average IP/INP levels of mossy cell markers. (D) Taqman analysis of mRNA levels of *Drd2* from Context+Food mice.

**Supplementary Figure 4. Validation of hippocampal *Drd2* function in mice.** (A) Immunohistochemistry and quantification of CamkIIa+/YFP+ cells in mice injected with AAVs expressing CamkIIa-mCherry and cre-dependent YFP in *Drd2*-cre mice. Scale, 200 *μ*m (B) Schematic drawing of experiment design (left), images showing c-fos staining (red) in *Drd2*-YFP (green) mice. Scale, 200 *μ*m Right panel shows quantification for c-fos+/yfp+ cells in the hippocampus. Unpaired Student’s t-Test, *p<0.05, n=3. (C) Images showing c-fos (green) staining in Drd2-mCherry (red) cells of corticosterone (530 nM) injected mice, vehicle, overnight water deprived mice and control mice. Quantification is shown in the right panels. Unpaired Student’s t-Test, n.s. Scale, 100 *μ*m (D) Left panel shows a schematic drawing of experiment design. Images showing the c-fos staining (red) in *Drd2*-YFP mice. Right panel shows the quantification of c-fos+/yfp+ cells. One-way ANOVA with Bonferroni correction, n=3. Scale, 200 *μ*m.

**Supplementary Figure 5. Immunohistochemical and electrophysiological analysis of DREADD-injected mice.** (A-C) Electrophysiology experiments to validate DREADD functionality in *Drd2*-cre mice. (A) Bright field image of neuron being recorded. White lines depict the patch clamp electrode. (B) mCherry fluorescence showing the viral expression in recorded neuron. White dashed lines depict the mCherry+ D2R neurons. (C and D) Baseline firing rate of DREADD-expressing *Drd2* neurons and firing rates in the presence of 10 μM of CNO. Unpaired Student’s t-test was used: *p<0.05. (E) Schematic drawing of experimental design. (F) mCherry (left) and c-fos (right) staining in AAV5-Camk2a-hM4Di-mCherry injected mice. Scale, 200 *μ*m. (G) mCherry (left) and c-fos (right) staining in AAV5-Camk2a-hM3DqmCherry injected mice. Scale, 200 *μ*m.(H) Cumulative food intake in saline injected mice expressing hM4Di virus. (I) Cumulative food intake in saline injected mice expressing hM3Dq virus. One-way ANOVA with Bonferroni correction was used for statistical analysis and no significant was found. (J) Scheme showing injection site and experimental design. (K) Food consumption (g) after optogenetic inhibition of Drd2 cells. (L) Food consumption (g) after optogenetic activation of Drd2 cells. Unpaired Student’s t-test, n=8, *p<0.05.

**Supplementary Figure 6. D2R cells activity is controlled by LEC CamkIIa+ cells.** (A) Schematic drawing showing the injection of monosynaptic tracing using cre-dependent TVAmCherry (red) and rabies virus (green) in the hippocampus. Scale, 310 *μ*m. Green cells depict neurons that contain a direct synaptic connection to *Drd2* cells. (B) ChR2-mcherry (red) staining in LEC and c-fos staining (green) after 5min of 10Hz stimulation with blue light (473 nm). (C) C-fos staining (green) in the hippocampus of ChR2-expressing and control LEC cells after 5min of 10Hz stimulation with blue light (473 nm). Mice/group=3. Scale, 200 *μ*m. (C) hM3Dq-expressing CamkIIa LEC cells were activated with CNO (1mg/kg) and c-fos staining (red) was observed in *Drd2*-YFP cells. White arrows show c-fos+ and *Drd2*+ cells. Inset shows mCherry-expressing CamkIIa cells, used as controls. Scale, 100 *μ*m.

**Supplementary Figure 7. Anxiolytic and locomotor behavior in Chr2- and Arch3.0injected mice.** (A) Open field test in *Drd2*-Arch3.0-expressing mice. (B) Open field test in *Drd2*-ChR2-expressing mice. Parameters analyzed: distance, velocity, time in center and time in border. Unpaired Student’s t-test, no significance was found, n=6.

**Supplementary Figure 8. Anxiolytic and locomotor behavior in DREADD-injected mice.** (A) Elevated plus maze test in *Drd2*-Dreadd-expressing mice. (B) Open field test in *Drd2*-DREADDs-expressing mice. One-Way ANOVA with Bonferroni correction, no significance was found, n=6.

## RELATED TO INFORMATION FOR TABLES AND VIDEOS

**Supplementary Table S1. RNA Sequencing data for Context+Food samples.**

**Supplementary Table S2. RNA Sequencing data for Context samples.**

